# Stress Recovery Triggers Rapid Transcriptional Reprogramming and Activation of Immunity in Plants

**DOI:** 10.1101/2023.02.27.530256

**Authors:** Natanella Illouz-Eliaz, Jingting Yu, Joseph Swift, Kathryn Lande, Bruce Jow, Za Khai Tuang, Travis Lee, Adi Yaaran, Rosa Gomez Castanon, Joseph R. Nery, Tatsuya Nobori, Yotam Zait, Saul Burdman, Joseph R. Ecker

## Abstract

All organisms experience stress as an inevitable part of life, from single-celled microorganisms to complex multicellular beings. The ability to recover from stress is a fundamental trait that determines the overall resilience of an organism, yet stress recovery is understudied. To begin unraveling the stress recovery process we studies recovery from drought stress in *Arabidopsis thaliana*. We performed a fine-scale time series of bulk RNA sequencing starting 15 minutes after rehydration following moderate drought. We reveal that drought recovery is a rapid process involving the activation of thousands of recovery-specific genes. To capture these rapid recovery responses in different leaf cell types, we performed single-nucleus transcriptome analysis at the onset of post-drought recovery, identifying a cell type-specific transcriptional state developing within 15 minutes of rehydration independently across cell types. Furthermore, we reveal a recovery-induced activation of the immune system that occurs independent of infection, which enhances pathogen resistance *in vivo* in *A. thaliana*, wild tomato (*Solanum pennellii)* and domesticated tomato (*Solanum lycopersicum* cv. M82). Since rehydration promotes microbial proliferation and thereby increases the risk of infection^1–2^, drought recovery-induced immunity may be crucial for plant survival in natural environments. These findings indicate that drought recovery coincides with a preventive defense response, unraveling the complex regulatory mechanisms that facilitate stress recovery in different plant cell types.

## Main

In most plants, extended periods of water deficit result in reduced growth, premature flowering, flower abortion, fruit abscission, and, ultimately, decreased yield^3–4^. Plant responses to drought have therefore been studied extensively as part of efforts to develop strategies or genetic manipulations that could mitigate the economic and agricultural consequences of future droughts.

Previous studies have identified and functionally characterized numerous genes that respond to water deficit, including various transcription factors (TFs) that regulate plant drought responses^4–6^. In general, drought stress in plants induces dramatic changes in the transcriptional landscape^7–8^. For example, the rapid up-regulation of genes involved in osmolyte accumulation enables water retention by adjusting cellular osmotic potential^9^. However, efforts to enhance drought tolerance through genetic manipulation have frequently resulted in undesired growth inhibition under non-drought conditions, thereby constraining the widespread engineering and adoption of drought-resilient crops^6,10–11^.

Given these limitations, we considered an alternative approach to investigating stress resilience in plants by focusing on post-drought recovery rather than drought resistance. Indeed, understanding a plant’s ability to recover from drought is central to a comprehensive understanding of drought resilience, as the potential for recovery defines whether the system can return to a stable state of function^12^. For example, drought recovery has been established as an indicator of drought tolerance in annual crops such as maize^13–14^, wheat^15–16^ and rice^17–18^. This extends to other abiotic stresses as well, for instance, the rates of submergence recovery in *Arabidopsis thaliana* (Arabidopsis) accessions were found to correlate with their submergence tolerance^19^. This finding suggests that the ability to recover after flooding is critical for plant survival and reproductive success. One of the well-characterized plant responses to drought alleviation is the downregulation of the drought responsive genes. For example, most drought-regulated genes recover to normal expression levels within three hours of rehydration^20^. However, Oono et al.^21^ identified 82 “recovery-specific” genes whose expression was drought-invariable but altered by subsequent rehydration.

Despite these known transcriptional responses during drought recovery, it remains unclear whether these responses constitute a conserved drought recovery mechanism activated throughout the entirety of plant cell types. Here, we explore the transcriptomic landscape in Arabidopsis throughout the drought recovery process using a high-resolution time series of RNA sequencing data. We identified over 3,000 recovery-specific genes that were differentially expressed between 15 minutes and 6 hours after rehydration. Using single-nucleus RNA-seq, we analyzed cell type-specific transcriptional signatures immediately after inducing drought recovery. We identified a recovery unique transcriptional signature common between sub-population of epidermal, hydathode, sieve element and mesophyll cells that is found only once drought recovery is initiated. Within the gene modules enriched in sub-populations we found many immune-associated genes. Based on these observations, we propose that some cells enter a recovery cell state upon rehydration. The results indicated that one of the first steps of the recovery process is the activation of a rapid preventive immune response. We therefore tested whether short-term recovery from moderate drought activated functional immunity to block pathogen proliferation in plant leaves. In agreement with this hypothesis, we found that drought recovery-induced activation of immune system genes leads to reduced disease severity and bacterial load in infected leaves of both Arabidopsis and two tomato species.

### Drought recovery-specific genes

To study the transcriptional response to drought recovery, we performed a fine-scale RNA-seq time series using vermiculite-grown Arabidopsis rosettes. We collected Arabidopsis rosettes during moderate drought and at seven subsequent recovery time points within the six hours following rehydration (Fig. a-b). At each time point, well-watered rosettes were also collected to control for diurnal transcriptional changes (Fig. 1b). We then conducted a differential expression (DE) analysis of the treatments and time points and defined two groups of genes: drought-responsive genes and recovery-specific genes. Drought-responsive genes were defined as genes that were DE compared to well-watered plants during the drought stress i.e., at time 0 (Extended Data Table 1). Conversely, recovery-specific genes were not DE during the drought period but were DE in any of the post-drought recovery time points (Extended Data Table 2).

**Figure 1.**
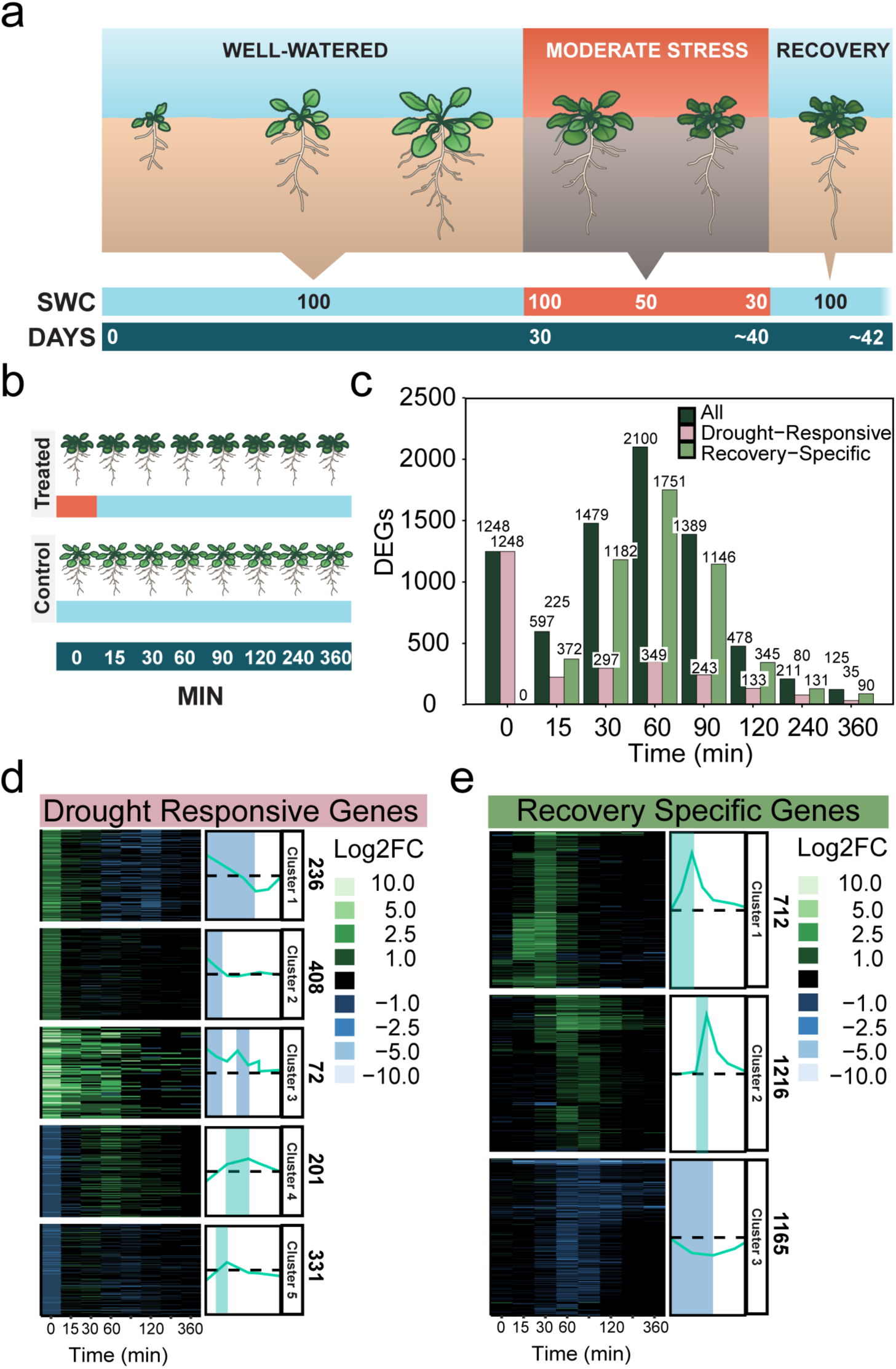
A fine-scale RNA-seq time course of drought recovery reveals recovery-specific genes. **a**, Illustration of the recovery time-course experimental design. Plants were grown in vermiculite under a short-day photoperiod. After 30 days, irrigation was stopped for drought-treated plants until they reached 30% relative soil water content (SWC). We then rehydrated the drought-treated plants by saturating the vermiculite to initiate drought recovery. We collected 3 biological samples from each time point of recovery, and well-watered control at each time point (n=3) **b,** Illustration of the samples collected for RNA-seq. We collected samples during moderate drought (t=0 at 30% SWC) and at seven additional time-points during the recovery process, from 15 min to six hours after rehydration. All samples from drought-treated plants were collected alongside equivalent samples from well-watered controls, with three replicates per treatment (drought or control) per time point. **c,** Number of differentially expressed genes (DEGs) at different time points during recovery. For each time point, we show the total number of DEGs as well as the numbers of drought-responsive and recovery-specific genes. DEGs were identified by comparing drought-treated samples to well-watered controls at each time point. **d-e,** K-means clustering and expression patterns of **d,** drought-responsive and **e,** recovery-specific genes.

Based on these definitions, we identified 1,248 drought-responsive genes, of which 662 (53%) were also DE during recovery. After only six hours of rehydration, 97% of these genes returned to normal expression levels (Fig. 1c-d). We additionally identified over 3,000 recovery-specific genes across all recovery time points (Fig. 1c and e), suggesting that drought recovery responses cannot be solely explained by the alleviation of drought induced transcriptional responses. We used K-means clustering to characterize gene expression patterns during drought and recovery. The drought-responsive genes exhibited five distinct expression patterns (Fig. 1d): (1) drought-induced but downregulated during recovery, (2) drought-induced but not DE during recovery, (3) drought- and recovery-induced; (4) drought-downregulated but recovery-induced; and (5) drought-downregulated but not DE during recovery. Recovery-specific genes exhibited three types of expression patterns based on whether they were induced in (1) early recovery or (2) late-recovery or (3) downregulated by rehydration (Fig. 1e). These results demonstrate that drought recovery involves the activation of thousands of recovery specific genes. These genes are timely regulated, with a set of genes being upregulated from as early as 15 mins after rehydration, and a later set of genes starting to accumulate after 30-60 mins.

### Global and cell type-specific recovery induced transcriptional reprogramming

To create study the leaf cell type-specific transcriptional responses upon drought recovery, we performed single-nucleus RNA-seq (snRNA-seq). For this experiment, we processed two replicates undergoing long-term moderate drought and an additional two replicates after 15 minutes of rehydration, as well as equivalent samples from well-watered controls. In this case, the well-watered controls were also provided with 15 minutes of additional irrigation to ensure that any changes in gene expression were specific to drought recovery and not due to mechanical root stress (Fig. 2a). After data quality control and filtering, our integrated dataset included over 144,000 single-nuclei transcriptomes. The median number of unique molecular identifiers (UMI) per nucleus was between 2,534 to 3,995 across the eight samples, and the median number of genes detected per nucleus was between 1,213 to 1,698 (average 1,530; see Extended Data Table 3).

**Figure 2.**
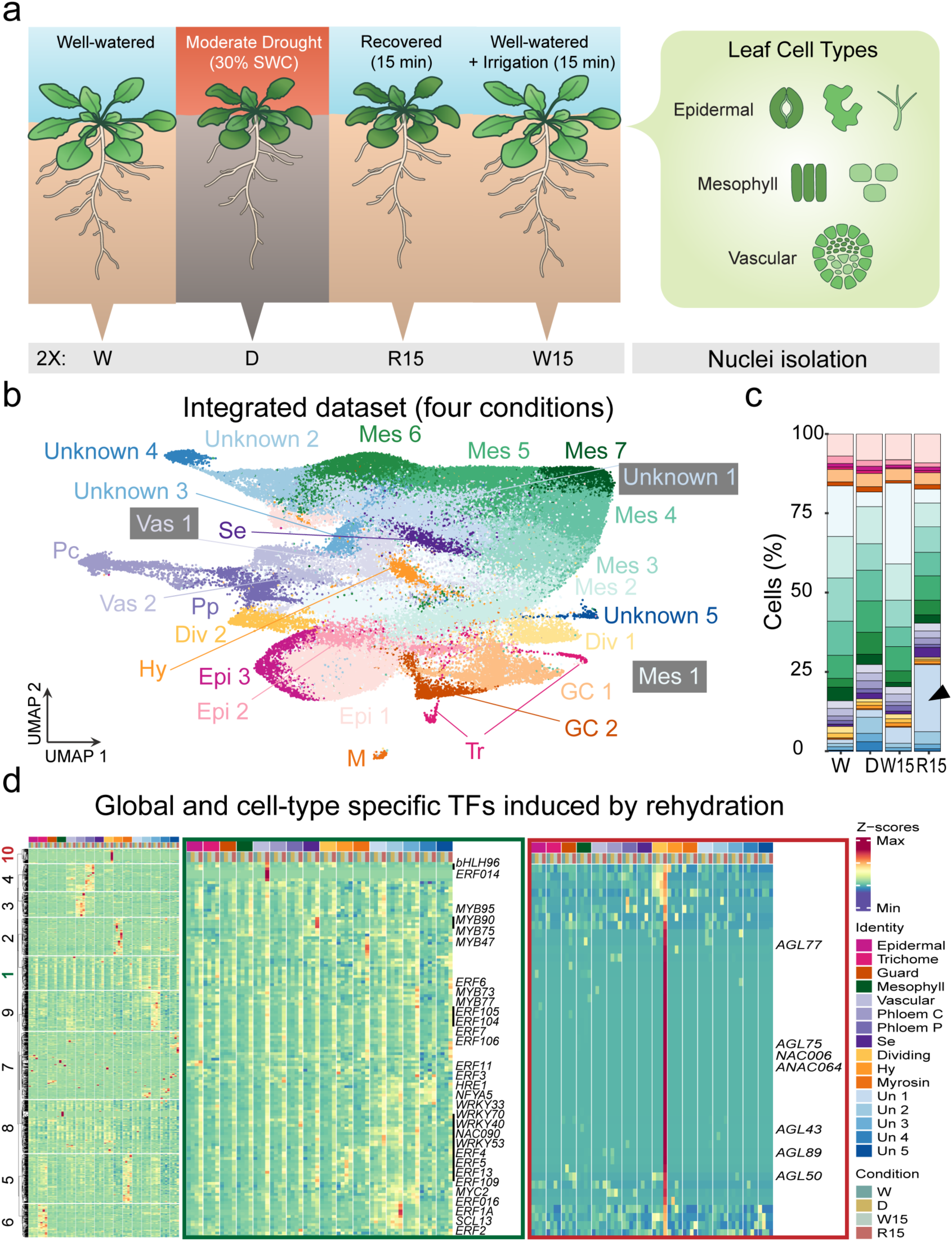
Drought recovery is initiated via transcriptional reprogramming in proliferating cells. **a**, Illustration of the experimental design used to generate a single-nucleus gene expression map. We collected two biological samples (n=2) from each of four conditions: (1) well-watered plants prior to drought [W]; (2) plants experiencing moderate drought [D]; (3) drought-treated plants after 15 min of rehydration [R15]; and (4) well-watered plants after 15 min of additional irrigation [W15]. **b,** Uniform Manifold Approximation and Projection (UMAP) of the integrated single-nucleus RNA-seq dataset revealed 27 cell identities encompassing the major leaf cell types: epidermal 1-3 (Epi), guard cell 1-2 (GC), trichome (Tr), hydathode (Hy), myrosin (M), mesophyll 1-7 (Mes), phloem parenchyma (Pp), phloem companion (Pc), sieve element (Se), and dividing 1-2 (Div). Five clusters could not be assigned to a cell type (Unknown 1-5). **c,** Cell identities recovered from each of the four conditions, expressed as a percent of the total number of recovered cells. Plants experiencing drought maintain similar cell identities, but the initiation of recovery leads to enrichment of one of the unidentified clusters (Unknown 1). **d,** Gene expression of Arabidopsis TFs during drought and immediate recovery represented by Z-score. In green square, cluster 1 genes, representing TFs globally induced by rehydration in most cell types. In red square, cluster 10 genes that were strongly induced by rehydration, specifically in dividing cells.

Unsupervised clustering of the snRNA-seq profiles resulted in 27 clusters representing unique molecular cell identities (Fig. 2b, Extended Data Figs. 1-2). All independent samples and treatment conditions integrated into the same clusters (Extended Data Fig. 1). We used three major practices to annotate cell identities for these clusters. First, we overlapped each cluster’s marker genes (genes uniquely or highly expressed in a certain cluster) with previously validated marker genes for all leaf cell types and tissues (Extended Data Tables 4-5 and Extended Data Figs. 3-4). Second, we projected our dataset on published single-cell datasets preformed on Arabidopsis leaves (Extended Data Fig. 5). Third, we manually investigated the top marker genes in each cluster and used Gene Ontology (GO) enrichment analysis to see enriched biological processes (Unknown clusters; Extended Data Fig. 6). Based on the genes expressed in each cluster and their expression levels, we were able to confidently assign cell types to 22 of the 27 clusters.

Once the clusters had been assigned to cell types, we investigated cell type-specific changes during the onset of drought recovery. A transcription factor (TF) expression analysis across the different treatments revealed that TFs are rapidly induced by rehydration across all cell types. These TFs had many genes from the *ETHYLENE RESPONSE FACTOR* (*ERF*) and *WRKY DNA-BINDING PROTEIN* (*WRKY*) families (Full list in: Extended Data Table 6, GO term enrichment analysis in: Extended Data Fig. 7). In contrast, some cell types, such as myrosin and vascular cells exhibited cell type-specific changes in TF expression upon recovery. An interesting group of TFs were rapidly activated in a contextual manner that was both treatment specific and cell state specific. These TFs were upregulated only in dividing cells at the onset of recovery (Fig. 2d). Four gene families dominated this set of genes: MADS box, B3, NO APICAL MERISTEM (NAC), and PAIRED AMPHIPATHIC HELIX (PAH2) families (Full list in: Extended Data Table 7). Based on a sequence similarity analysis, the highly activated genes within each family were phylogenetically clustered. For example, the upregulated MADS box genes were all type I MADS box (M-type) genes: *AGAMOUS-LIKE43* (*AGL43*), *AGL89*, *AGL77*, *AGL75*, *AGL74*, *AGL84*, and *AGL50*.

Similarly, all the B3 TFs induced by drought recovery in dividing cells were in the REPRODUCTIVE MERISTEM (REM) branch^22^, despite the B3 family containing multiple distinct gene subfamilies^23^. These findings suggest that drought recovery triggers cell type-specific transcriptional reprogramming.

### Recovery cell states in distinct cell types

To study heterogeneity within cell types during drought recovery, we further subclustered cells within each cell type based on their gene expression profiles (Extended Data Figs. 8-9). In epidermal, mesophyll, hydathode, and sieve-element cells, some of the cell subpopulations were enriched (i.e., over 50% of cells in the subcluster) with cells from the drought recovery treated plants (Fig. 3a, Extended Data Fig. 9). We hypothesized that these cell subpopulations form distinct subclusters due to their transition into a recovery cell state (RcS). To test this hypothesis, we performed a motif enrichment analysis on each of the suspected RcS subclusters to determine whether we could identify regulatory motifs that were common to the putative RcS across all cell types. We found that all the subclusters that we identified as having an activated RcS were enriched with the *CALMODULIN BINDING TRANSCRIPTION ACTIVATOR 1* (*CAMTA1*) and *CAMTA5* binding motifs (Fig. 3b). In Arabidopsis, the CAMTA family of TFs regulate plant defense and stress responses by modulating the expression of genes involved in pathogen defense, abiotic stress tolerance, and plant development. These TFs interact with calcium-bound calmodulin to mediate downstream signaling pathways^24^. To investigate if CAMTAs specifically regulate the RcS, we performed motif enrichment analysis on all subclusters to see if the *CAMTA1* and *CAMTA5* motifs were enriched in other subclusters. Out of 162 subclusters, only 9 subclusters were enriched with *CAMTA* motifs (Extended Data Table 8, and Extended Data Fig. 10), suggesting that *CAMTA1* and *CAMTA5* are nearly exclusively enriched in RcS subclusters, and could be the regulators of the formation of this cell state.

**Figure 3.**
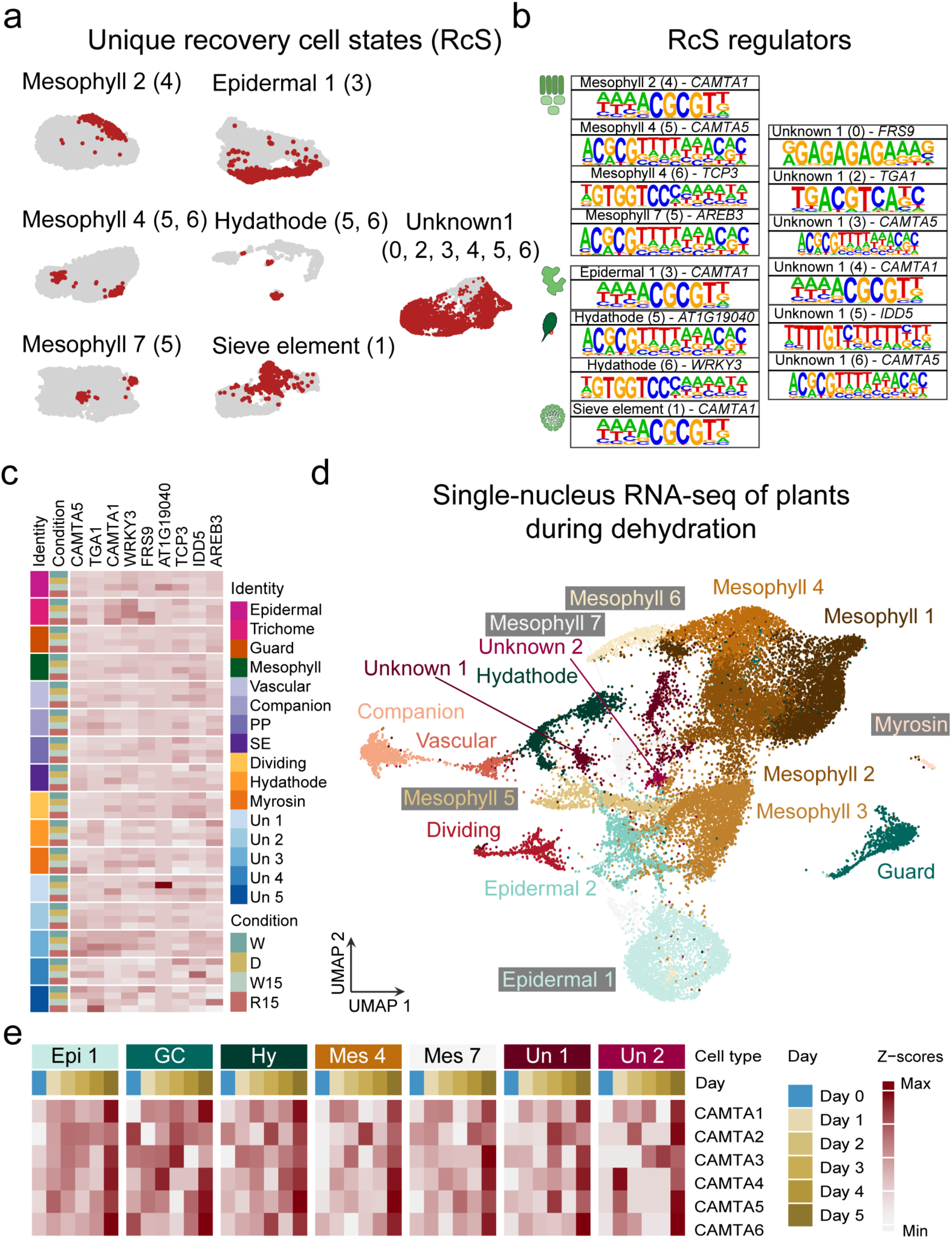
Recovery cell states (RcS) appear in sub-populations of epidermal, mesophyll, hydathode, and vascular cells. **a**, Subclustering of cells within each cell type from Fig. 2C revealed unique subclusters enriched in cells from plants recovering from drought. Drought-recovered cells are indicated in red. **b,** Putative regulators of the RcS, identified using *de novo* motif enrichment analysis performed for each recovery-enriched subcluster. The gene name above each motif is the predicted TF that binds the enriched motif. **c,** Z-score representation showing the expression levels of the TFs that putatively regulate the formation of the post-drought RcS. **d,** Uniform Manifold Projection and Approximation (UMAP) of the snRNA-seq data generated from *Arabidopsis* rosettes over the course of a five-day dehydration experiment. We collected multiple rosettes for each day and processed them as one biological replicate (n=1). **e,** Z-score representation showing the expression levels of predicted CAMTA TFs during the early stages of plant dehydration.

We predicted that *CAMTA1* and *CAMTA5* regulate the RcS; however, neither the CAMTA genes nor the cell type-specific TFs predicted to regulate the RcS were upregulated in response to rehydration (Fig. 3c). Since the RcS is rapidly activated, we hypothesized that *CAMTA* transcripts may be differentially expressed during the early stages of drought and would therefore not be observed in our data, which was collected after moderate drought had established. To test this hypothesis, we performed an additional snRNA-seq experiment on Arabidopsis plants harvested each day over the course of five days starting from the onset of drought (Fig. 3d, Extended Data Fig. 11). Cell identities were identified using unsupervised clustering, as before. We then investigated the expression of *CAMTA* genes in different cell types during early drought. In most cell types, *CAMTA* genes were upregulated in the days following dehydration (Fig. 3e, Extended Data Fig. 12), suggesting that the regulation of *CAMTA* is very sensitive to changes in water availability and that *CAMTA* transcripts accumulate early in response to drought.

In addition to testing for a common regulatory pathway, we investigated whether the RcS subpopulations within the different cell types expressed similar gene networks by performing gene co-expression network analysis^25^ for the seven cell clusters with an activated RcS subpopulation in the drought recovery experiment (Fig. 3a). For each cluster, this analysis identified gene modules (Extended Data Fig. 13-14). We next searched for common hub genes (Genes with high module membership (kME) and high intramodular connectivity (kIN)) among the modules that were enriched in RcS cells from different cell types to identify genes that have common key roles in the formation of recovery cell states or its function. Overall, we found 212 hub genes, of which 50 were shared by at least two modules (Extended Data Table 9). Interestingly, in the immediate recovery snRNAseq, 70% of the top marker genes for cluster “unknown 1” overlap with the hub genes identified in the RcS gene network. This finding led us to hypothesize that cluster “unknown 1” represents a cell state that is superimposed across different cell types at the onset of drought recovery (Extended Data Fig. 15).

Overall, the suite of common hub genes among the six RcS subclusters from different cell types suggest that the functional onset of drought recovery in RcS subpopulations is comprised of cell wall modifications and the regulation of nutrient uptake, as well as cytoplasmic detoxification processes and DNA repair. The most prevalent hub genes shared among RcS subclusters are genes that play a critical role in cell growth and plant development; these genes included *XYLOGLUCAN ENDOTRANSGLUCOSYLASE/HYDROLASE 22* (*XTH22*)^26^, *SLAC1 HOMOLOGUE 3* (*SLAH3*)^27^, and *EXORDIUM-LIKE1* (*EXL1*)^28^. Since toxins accumulate in plant cells during drought stress^29^, we suspected that detoxification may also be an important part of recovery. Indeed, our cross-cluster hub gene analysis implicated *ARABIDOPSIS THALIANA DETOXIFICATION 1* (*AtDTX1*), which is localized in the plasma membrane of plant cells and mediates the efflux of plant-derived or exogenous toxic compounds from the cytoplasm^30^. An additional hub gene common across RcS cell populations was *CHROMATIN ASSEMBLY FACTOR 1* (*CAF-1*) *AtCAF1a*, which facilitates the incorporation of histones H3 and H4 onto newly synthesized DNA. The absence of the CAF-1 chaperone complex results in mitotic chromosome abnormalities and changes in the expression profiles of genes involved in DNA repair^31^. In addition, AtCAF1 proteins regulate mRNA deadenylation and defenses against pathogen infections^32^. Taken together, these finding suggest that drought recovery triggers the formation of a specific cell state imposed within several leaf cell types.

To identify more general processes involved in drought recovery, we performed a cell type-specific differential expression analysis comparing plants harvested 15 minutes after the onset of drought recovery to (i) plants experiencing drought and (ii) the well-watered controls that received additional irrigation for 15 min. Both controls were used for DE analysis to ensure that DE genes were not simply maintained from their changes during drought or induced by the mechanical stress of additional watering. In nearly all cell types, photosynthesis and carbon fixation were suppressed during the onset of drought recovery. The only exception was in one of the seven mesophyll cell clusters, which maintained photosynthetic activity but downregulated the expression of genes associated with bacterial responses (Extended Data Fig. 16a). Conversely, genes that were upregulated at the onset of recovery were associated with diverse stressors including wounding, cold, heat, light, and reactive oxygen species (i.e., oxidative stress). The upregulation of such a broad suite of stress response genes suggests that the plant perceives the onset of recovery as a state of stress, perhaps because recovery presents a change in the current homeostasis that was established during drought.

### Recovery-induced immune responses

We found evidence for the activation of multiple immunity-related genes in the top shared hub genes among RcS subclusters. Among these genes for example, *TETRASPANIN 8* (*TET8*) has been shown to be upregulated in response to the Flg22 and Elongation Factor Tu (EF-Tu) elicitors, which mimic pathogen infection^33^. Another shared hub gene *SALT-INDUCIBLE ZINC FINGER 1* (*SZF1*) also regulates plant immunity, as *szf1,2* knock-out mutants show increased susceptibility to *Pseudomonas syringae* pv. *tomato* DC3000 (*Pst* DC3000)^34^. Three other shared hub genes (*AT5G41750, AT5G41740* and *AT4G19520*) encode TOLL INTERLEUKIN RECEPTOR (TIR)–type NB-LRR proteins and thus belong to the most common class of disease resistance genes in plants^35^. Another example is *ACTIVATED DISEASE RESISTANCE 2* (*ADR2*) that enhances resistance to biotrophic pathogens^36^. Finally, we also found *EARLY RESPONSIVE TO DEHYDRATION 15* (*ERD15*) as a top shared hub genes within the RcS enriched modules, which belongs to a small and highly conserved protein family that is ubiquitous but specific to the plant kingdom. Overexpression of ERD15 proteins in response to various pathogen elicitors was shown to improve resistance to known pathogens and has also been shown to impair drought tolerance^37^. Thus, the activation of a gene such as *ERD15* specifically upon rehydration maybe beneficial to contribute to plant defense without compromising the drought response.

Other genes upregulated during the recovery initiation were implicated in processes associated with plant immunity, including genes involved in response to bacterial pathogens, immune system processes, responses to jasmonic acid (JA), and cell death. We found additional processes associated with plant defense that were upregulated in specific cell types; for example, genes involved in the responses to oomycetes or ethylene response were upregulated in almost all cell types during recovery, whereas defense responses against insects were only upregulated in one-third of epidermal cell clusters, 50% of vascular cell clusters, and two of the “unknown” clusters (Extended Data Fig. 16b).

We suspected that the transcriptomic upregulation of immune system processes during drought recovery is an innate response to post-drought rehydration rather than a proactive response to microbial infection following rehydration. To test this hypothesis, we examined the upregulation of recovery induced genes under axenic conditions. Arabidopsis seedlings were grown on sterile agar plates for 14 days and then transferred to low-water-content agar plates^38^, which we produced by decreasing the water content of the media to 50% (thus imposing moderate osmotic stress) (Extended Data Fig. 17a-b). After 14 days on the low-water-content plates, plants were rehydrated and collected at different recovery time points (0, 15, 30, 90 and 120 min) for bulk RNA-seq analysis. The upregulation of the known drought marker genes *RESPONSIVE TO DESICCATION 29A (RD29A), RD20*, and *DELTA1-PYRROLINE-5-CARBOXYLATE SYNTHASE 1 (P5CS)*^39–41^ confirmed that the plants were experiencing moderate drought stress (Extended Data Fig. 17c). We additionally found that immune-related genes were upregulated after 15 minutes of rehydration even in the axenic system (Extended Data Fig. 17d). These results support the hypothesis that plants activate a prophylactic defense response upon recovery from moderate drought, independent of pathogens in their environment.

Given the strength of this immunity-related transcriptional response during drought recovery, we used data from Bjornson et al.^42^ to evaluate how the genes implicated in our analysis correlated with known transcriptional responses to biotic elicitors. Bjornson et al.^42^ performed transcriptomic analysis across a fine-scale time series to study rapid signaling transcriptional outputs induced by well-characterized elicitors of pattern-triggered-immunity (PTI) in Arabidopsis. Their results showed that the transcriptional responses to diverse microbial- or plant-derived molecular patterns are highly conserved. When we aligned these identified transcriptional responses with the recovery specific genes identified by the bulk RNA-seq data from the first two hours of our drought recovery time series (Fig. 1), we found that 50% of recovery-specific genes upregulated after 15 minutes of rehydration overlap with the core responses to biotic elicitors in the Bjornson et al.^42^ dataset. More generally, this analysis shows a rapid and robust peak of immune-relevant gene expression that gradually decreased with time since rehydration (Fig. 4a).

**Figure 4.**
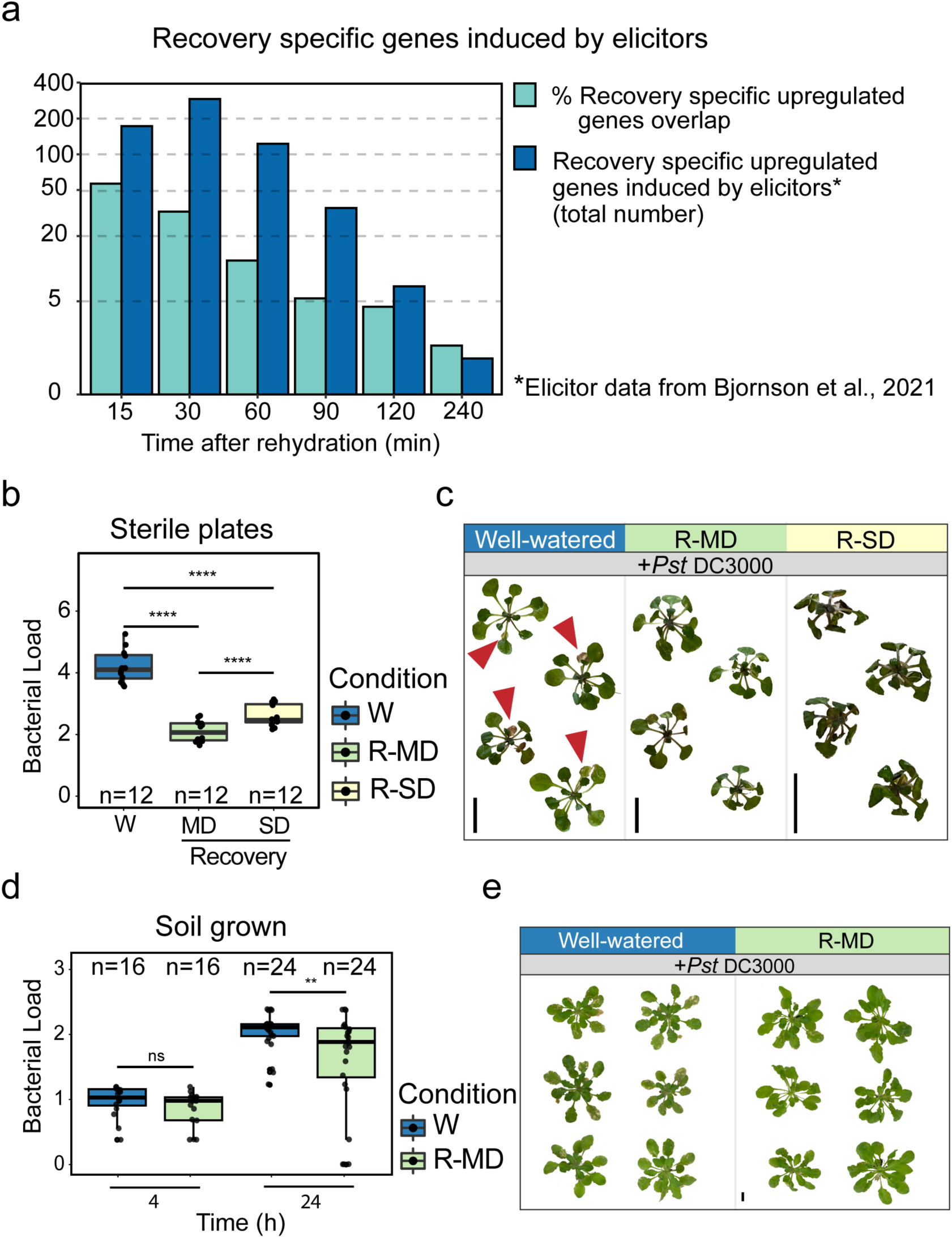
Recovery from moderate drought enhanced pathogen resistance in *Arabidopsis*. **a**, Overlap between upregulated recovery-specific genes in our bulk RNA-seq data and genes upregulated by different biotic elicitors as reported by Bjornson et al. (2021). For each recovery time point, bars show both the percentage of upregulated recovery-specific genes that are also known to be activated by biotic elicitors as well as the total number. **b,** Bacterial load (log_10_CFU) of plants that were grown on sterile low-water-content plates and then submerged in a suspension of *Pseduomonas syringae* pv. *tomato* DC3000 (*Pst* DC3000, OD_600_=0.005) or a control solution. Bacterial growth was measured two days post-inoculation, and well-watered controls (W) were compared against plants recovering from either moderate (R-MD) or severe drought (R-SD). For this analysis, we merged two independent experiments for a total of n=12 per treatment. Significance values were calculated with a two-way ANOVA of treatment and batch followed by Tukey’s *post hoc* test. P-values were FDR-corrected. **c,** Representative images of plants taken four days post-inoculation. **d,** Bacterial load (log_10_CFU) of soil-grown plants sprayed with *Pst* DC3000 (OD_600_=0.05) or a control solution after experiencing moderate drought. Bacterial growth was measured 4 and 24 hours post-inoculation and compared against controls using a two-way student’s t-test. We collected 16 leaves (n=16) to measure bacterial load at the initial time point, and 24 leaves (n=24) to measure bacterial growth 24 hours post infection. **e,** Representative images of plants taken 14 days post-inoculation. In **b** and **d**: ns = P > 0.05, * = P ≤ 0.05, ** = P ≤ 0.01, *** = P ≤ 0.001, **** = P ≤ 0.0001. Boxplots middle line shows the median, the lower and upper hinges are the 25th and 75th percentile, respectively. The whiskers extend from the hinges to the most distant value within 1.5 * IQR of the hinge, where IQR is the inter-quartile range, or distance between the first and third quartiles.

### Drought recovery-induced immunity *in vivo*

To test the functionality of post-recovery immune activation, we examined whether the activation of immunity-related genes during short-term recovery from moderate drought promotes pathogen resistance *in vivo*. We used the sterile low-water-content plate system and examined recovery from both moderate (40% water-content) and severe stress (25% water-content), as described above. Plants from the control and moderate or severe osmotic stress conditions were rehydrated for 90 minutes and then inoculated with *Pst* DC3000 via submersion in a bacterial suspension. Whole rosettes were collected and weighed 48 hours following infection and used to quantify bacterial growth. Plants recovering from drought had significantly lower bacterial concentrations than control plants, and recovery from moderate stress suppressed bacterial growth better than recovery from severe stress (Fig. 4b-c). These results suggest that recovery from moderate drought enhances resistance to *Pst* DC3000.

To further validate these results, we grew plants on soil (non-sterile conditions) for 30 days under short-day conditions. Drought-treated plants were transferred to dry trays and dehydrated for one week down to 30% relative soil water content, while control plants continued to receive regular irrigation. We rehydrated drought-treated plants for 90 minutes and then sprayed the leaves of both control and recovered plants with *Pst* DC3000. Leaf discs were collected from each treatment 4 and 24 hours after inoculation, surface sterilized, and then used to measure bacterial load. The bacterial load 4 hours after inoculation did not differ between well-watered and drought-recovered plants. However, 24 hours after inoculation, the bacterial load of drought-recovered plants was significantly lower than that of controls, indicating increased resistance to *Pst* DC3000 in rehydrated plants (Fig. 4d-e). We call this response, which was consistent across our two separate empirical tests, drought recovery-induced immunity (DRII).

Given the consistency of the DRII response in Arabidopsis, we further tested whether DRII is conserved among various plant species and in response to different pathogens. First, we examined pathogen proliferation in wild tomato plants recovering from moderate drought and infected with either *Pst* DC3000 or an additional tomato pathogen *Xanthomonas perforans* strain 97-2, which causes the tomato spot disease^43^. For this test, we used the drought-tolerant tomato species *Solanum pennellii*^44^. We grew *S. pennellii* plants at 25°C with a photoperiod of 12 hours light and 12 hours dark. When the plants had two to three true leaves (∼4 weeks old), we exposed them to moderate drought by stopping irrigation until the soil reached 30% water content relative to saturated pots, as we had done for Arabidopsis. Drought was then alleviated by irrigating pots to full saturation, and plants were infected 90 minutes after rehydration. Both the drought-recovered plants and well-watered controls were syringe-infiltrated with a suspension of either *X. perforans* 97-2 or *Pst* DC3000.

We first assessed disease severity in the well-watered and drought-recovered infected leaves by imaging inoculated leaves 5 days post infection and calculating the percentage of symptomatic area relative to the entire leaf surface. Leaves inoculated 90 minutes after recovery from moderate drought were significantly less symptomatic than leaves from well-watered controls (Fig. 5a-b). To directly measure bacterial concentrations in leaves, we compared leaf samples at an initial time point 4 hours after inoculation to samples collected 5 days post-inoculation. Although a similar number of bacteria entered the leaves of all plants, the plants that recovered from drought had a significantly lower bacterial load after five days (Fig. 5c). Our results were consistent in both tested pathogens: regardless of the pathogen, tomato leaves that recovered from drought exhibited lower disease severity and reduced bacterial concentrations than control plants. Our observation that short-term recovery from moderate drought increases pathogen resistance across all treatments further suggests that DRII is a preventive immune response that enhances pathogen resistance during the initial period of drought recovery.

**Figure 5.**
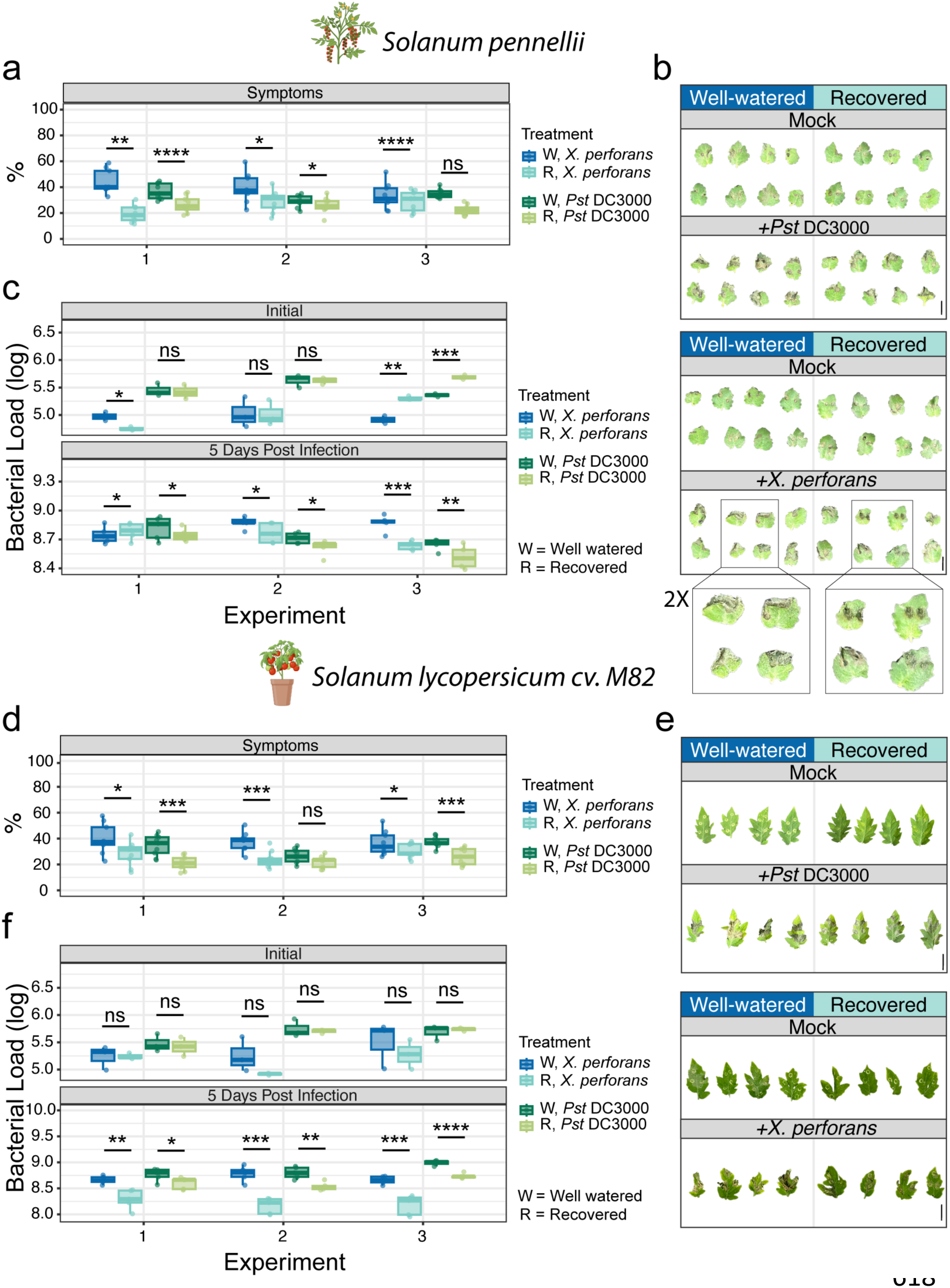
Drought recovery-induced immunity enhanced resistance to *X. perforans* and *Pst* DC3000 in wild (*Solanum pennellii*) and domesticated (*S. lycopersicum* cv. M82) tomato species. **a**, Lesion percentage analysis of *S. pennellii* leaves five days after being syringe infiltrated with either *X. perforans* OD_600_= 0.02 or *Pst* DC3000 OD_600_= 0.02. Inoculation was performed after 90 mins of recovery from moderate drought, and drought-treated samples were compared to well-watered controls. Results are shown for three independent experiments, 9 leaves were analyzed in each experiment (n=9). **b,** Representative images of well-watered or drought-recovered *S. pennellii* leaves infected with one of *Pst* DC3000, *X. perforans*, or a mock solution. Scale bar = 1 cm. **c,** Bacterial load (log_10_CFU) of the *S. pennellii* leaves measured five days post-infection. Results are shown for three independent experiments. 2 leaf discs (0.5 cm in diameter) were prepared for each sample, 5 samples were used for each independent experiment (n=5). **d,** Lesion percentage analysis of M82 leaves after undergoing the same post-drought infection experiment described for *S. pennelli*. 9 leaves were analyzed in each experiment (n=9). **e,** Representative images of M82 leaves infected with *Pst* DC3000, *X. perforans*, or a mock solution. Scale bar = 2 cm. **f,** Bacterial load (log_10_CFU) of M82 leaves measured five days post-infection. Results are shown for three independent experiments. 2 leaf discs (0.5 cm in diameter) were prepared for each sample, 5 samples were used for each independent experiment (n=5). Significant differences in all panels were identified using student’s t-test (ns = P > 0.05, * = P ≤ 0.05, ** = P ≤ 0.01, *** = P ≤ 0.001, **** = P ≤ 0.0001). Boxplots middle line shows the median, the lower and upper hinges are the 25th and 75th percentile, respectively. The whiskers extend from the hinges to the most distant value within 1.5 * IQR of the hinge, where IQR is the inter-quartile range, or distance between the first and third quartiles.

To further investigate DRII in an agriculturally relevant plant species, we repeated our post-drought immune challenge experiment using a cultivar of domesticated tomato, *Solanum lycopersicum* cv. M82 (M82), and the same two pathogens as before (*Pst* DC3000 and *X. perforans* 97-2). Because breeders focus on traits such as fruit size, shelf life, and brix, among others, we suspected that the DRII response could be lost during domestication. However, M82 plants showed similar results to *S. pennellii* upon infection with either pathogen (Fig. 6d-f), indicating that the DRII response we observed in Arabidopsis and *S. pennellii* following short-term recovery from moderate drought is retained in domesticated tomato. We think that the robustness of the DRII trait to the domestication processes underscores its potential significance in agricultural contexts.

## Conclusion

As large regions of the world gradually become drier, drought has become one of the most pressing threats to global food security^45–46^. Plant drought responses have therefore been studied extensively over the past decades, but the molecular mechanisms for drought recovery have been largely overlooked. We advanced the understanding of drought recovery processes by using bulk RNA-seq over a six-hour time series to identify recovery-specific genes. The resulting data enriches the pool of potential targets for developing drought-tolerant crops. For example, the genes whose expression was drought-invariable but downregulated during recovery (“cluster 3” in Fig. 1e) could be viable candidates for mutation screens aimed at improving post-drought effects and recovery dynamics without affecting drought resistance. Furthermore, the construction of a single-cell RNA-seq-based cell atlas of plants recovering from moderate drought allowed us to explore cell type-specific transcriptional changes in response to drought and immediate recovery. We were able to identify the major cell types in our dataset, and our findings suggest that the five unidentified clusters (Fig. 2b) most likely contain multiple cell types that clustered together because they were engaging in similar molecular or biochemical process during drought recovery, and therefore cluster by the transcriptional signature of a specific cellular state. As plants must constantly adapt to environmental changes, the molecular phenotypes of individual cells might vary to facilitate the metabolic and biochemical changes essential for adaptation to the current stressor.

Using our drought recovery single cell atlas, we identified the first regulatory steps underlying the transcriptional plasticity that allows plants to restore normal growth following drought. One of these steps involves upregulating a set of TFs specifically in dividing cells within 15 minutes of rehydration. However, little is known about the TFs identified by this analysis, although they largely belonged to four known TF families—the MADS box, B3, NAC, and PAH2 families. The MADS box family of TFs are well known for regulating flowering^47–48^. However, *AGL16* was shown to negatively regulate drought resistance via its effects on stomata^49^. The B3 family members contains a highly conserved DNA-binding domain and are involved in a variety of biological processes, many of which are related to the regulation of flowering through polycomb silencing^50–52^. The NAC TFs play important roles in development as well as abiotic and biotic stress responses^53^. Overall, the within-family phylogenetic proximity of the specific TFs upregulated by recovery suggest that these TFs act together to play a functional role in drought recovery, though determining the specific role of each TF will require further investigation of higher-order mutants.

Recently, Oliva et al.^55^ characterized a highly active transcriptional state that affects a subset of cells in different cell types across the developmental axis of the Arabidopsis root tip. The conserved cell state was named environmentally responsive state (ERS), since many ERS enriched genes were previously identified as responsive to various environmental cues. Here, we identified a transcriptional recovery cell state (RcS) that is triggered by environmental stimuli. This state emerges in different cell types rapidly and independently. We hypothesize that the RcS formation is governed by a small set of TFs, some of which are common among cell types such as *CAMTA1* and *CAMTA5*. It is possible that some cell type-specific TFs are also involved in the regulation of RcS depending on the cell type that the state was imposed on.

Our findings also show that transcripts for the calcium dependent *CAMTA* TFs accumulate during early drought, and that, although the upregulation of *CAMTAs* does not persist during drought recovery, many of the genes abundant during the RcS are regulated by *CAMTA1* and *CAMTA5*. Previous studies have shown that calcium flux can be affected by changes in water availability (both water deficit and water uptake)^54^. We therefore hypothesize that *CAMTA1 and CAMTA5* expression is induced during the early stages of soil dehydration and that CAMTA proteins are post-translationally stabilized. CAMTAs activate gene expression by interacting with calcium-bound calmodulin (CaM), and the influx of calcium upon rehydration leads to an increase in calcium-bound calmodulin that then activates CAMTA proteins to bind DNA and hence regulate the subsequent rehydration response. CaM levels may therefore be a limiting factor in the regulation of the rapid CAMTA1-dependent gene activation that we observed in the initial stages of drought recovery.

Our analysis suggest that the onset of drought recovery is coupled with the activation of a preventive immune response. Plants have evolved complex defense networks in response to microbial pathogens^35^. Aerial pathogens like *Pst* DC3000 or *X. perforans* mainly enter plants through stomatal pores, but the stomata close in response to drought stress^56^, making it harder for shoot pathogens to successfully attack plants experiencing drought. It should be noted that some pathogenic bacteria have evolved specific virulence factors to cause stomatal opening^57^. Moreover, plant immune responses can be sporadically induced by the circadian clock even in the absence of a pathogenic threat, a process driven largely by daily oscillations in humidity^58–59^. Such responses allow plants to prepare for the increased risk of infection when microbes are anticipated to be most infectious. Because rehydration promotes both pathogen proliferation and stomatal opening at a time when the immune system has already been suppressed by drought, plants are particularly vulnerable to pathogen attack during the initial stages of drought recovery^60–61^. The rapid upregulation of immunity genes that we observed in our study may therefore be crucial for ensuring plant survival in water-fluctuating environments. We accordingly propose that DRII is a preventive immune response that evolved to confer resistance against pathogens during drought recovery.

More broadly, our research shifts the focus from the well-studied drought responses to plant transcriptional activity during the recovery period. Although plant responses to drought have been extensively studied for over a century, these findings have yet to be successfully translated to the development of drought-tolerant crops^62^. Our results highlight the presence and importance of recovery-specific mechanisms that could be targeted in future experiments and engineering projects. We suggest a new avenue for research in which efforts to improve crop resilience should focus not only on survival during drought but also on rapid and effective recovery. By targeting recovery-specific genes, we can now endeavor to develop crops that not only withstand drought but also recover more swiftly, ensuring minimal yield loss and sustainable food production in an era of increasing climate unpredictability.

## Methods

### Plant materials and growth conditions

*Arabidopsis thaliana* Columbia-0 (Col-0) background was used throughout this study. Col-0 seeds were sterilized with chlorine fumes generated by mixing 100 ml bleach and 4 ml hydrochloric acid (HCl 1M). Sterilized seeds were sown on large square petri dishes with ¾ Linsmaier & Skoog with buffer (LS) media per liter and 2% agar, and then stratified at 4 °C for three days. Plates were placed in a growth chamber and grown in short-day conditions (8h light: 16h dark) at 22 °C with a light intensity of ∼110–130 μmol m^−2^ s^−1^. For drought treatments, seedlings were transferred to vermiculite pots two weeks after germination on plates, with 6 seedlings per pot. Tray size was 27.79 cm width, 54.46 cm in length, and 6.2 cm depth. Vermiculite was saturated with ¾ LS liquid media before the seedling transfer, 2L media per tray. After two days, each tray was watered with 2L ultra-pure water, and this continued for two weeks until the drought treatment started. When plants were 30 days old, each pot was weighed and transferred to a dry tray. The pots in the tray were weighed daily, and relative water content was calculated. The experiment started when the pots reached 30% relative soil-water content (SWC). Three whole Arabidopsis rosettes per treatment were collected separately as an independent sample, time 0 was 9 am (one hour after the lights turn on in the chambers). Three samples per treatment were collected alongside a well-watered control at each time point. For sterile low-water agar experiments, plates were prepared modified from Gonzales et al., 2023; 100% - 4L DDW, 3 LS bags, 80g agar – 120ml per plate, 50% (moderate stress) - 2L DDW, 3 LS bags, 80g agar – 60ml per plate, and 25% (severe stress) - 1L DDW, 3 LS bags, 80g agar – 30ml per plate.

For the drought time course experiment, plants were grown on plates and transferred to vermiculite as described above with the notable exception that transfer from plates took place 17 days after germination and remained on saturated vermiculite for 12 days before drought initiation. Drought was initiated by draining excess media from the pot and equilibrating to 100% relative SWC. Whole rosettes (n= 3 – 6) were sampled and frozen 4.5 hours after subjective dawn each day for 5 days.

Col-0 seeds were sterilized and seeded on plates. Plates were kept in the dark at 4C for three days. After three days, plates were moved to a growth chamber with short-day photoperiod light and 22 °C temperature.

Tomato seeds of *Solanum pennellii* and *Solanum lycopersicum* cv. M82 were germinated in a petri dish with 1/2 Murashige & Skoog (MS) with vitamins and FeNaEDTA (Cat# 07190008) media (no sucrose added). Ten-days-old seedlings were transferred to tray pots containing a mixture of vermiculite (Agrekal Moshav Habonim) and commercial soil (Tuff A.C.S) (1:1 by volume) and grown in a 12-hour light-dark cycle (12h light: 12h dark) at 25°C.

### Bacterial inoculation

*Pseudomonas syringae* pv*. tomato (Pst)* DC3000 was grown overnight in 10 ml liquid King’s B medium containing rifampicin (40 µg/ml) (Cat#: R3501-250MG, MilliporeSigma, MA) and tetracyclin (10 mg/ml) (Cat#: T3258, MilliporeSigma, MA) at 28°C for 24 h. Before adjusting the density, bacterial cells were washed two times with autoclaved water followed by centrifugation at 6,000 rpm for 2 min and re-suspension in water. For bacterial growth assays, well-watered and 90 min recovered Arabidopsis plants were inoculated with *Pst* DC3000 (OD_600_ = 0.005-0.5, depending on the experiment and inoculation method). For spray treatment Silwett77 (Cat#: S7777, PhytoTachLabs, KS) was added to bacteria before spraying. Bacterial titer was assessed as the log_10_ transformed colony forming units (CFU) per plant weight when collecting whole plants from sterile plates.

For experiments performed on tomato, syringe infiltration was used for bacterial inoculation. For bacterium counting, ten leaf discs (0.5 cm in diameter) were prepared from the second and third true leaves using a hole puncher (two discs per leaf) for each sample, five samples were used for each independent experiment. Each sample was separately homogenized in 10 mM MgCl_2_. The homogenate was serially diluted in 10 mM MgCl_2_. 10μl of diluted homogenate were plated on antibiotics and LB agar media on a square petri dish and incubated at 28°C for 48 h for determination of bacterial concentrations in leaves (log^10^ CFU/cm^2^).

### Assessment of disease severity in in tomato leaves

Disease severity in tomato (*S. pennellii* and *S. lycopersicum* cv. M82) leaves was assessed by calculating the percentage of the symptomatic area in the leaves relative to the whole leaf area. The calculation was done based on threshold color and color space HSB in ImageJ version 1.54d, according to Tuang et al.^1^ Nine leaves per experiment were analyzed for the lesion percentage (%).

### RNA extraction, bulk RNA library construction

Total RNA was extracted from three independent biological replicates of each time point using RNeasy Plant Mini Kits (Cat#79254, Qiagen, CA). Tape Station checked RNA quantity for quality control. Library construction was performed using Illumina Stranded mRNA Prep (Cat#20040534, Illumina, CA).

### Nuclei extraction and single-nuclei library construction

Seedlings were transferred to vermiculite pots two weeks after germination in trays, 6 seedlings per pot. Vermiculite was saturated with ¾ LS liquid media before the seedling transfer, 2L media per tray. After two days each tray was watered with 2L ultra-pure water, and this continued for two weeks until the drought treatment started. When plants were 30 days old, each pot was weighed and transferred to a dry tray. The experiment began when each pot reached 30% SWC. Between 12-18 whole rosettes were collected for each time point * condition sample. We used mortar and pestle to grind the frozen tissue. Powdered tissue was then placed in a nuclei extraction buffer [NEB; 500ul 1M TRIS pH=7.4 (Cat# 15567027, Thermo Fisher Scientific, MA), 150µl 1M MgCl_2_ (Cat# AM9530G, Fisher Scientific, MA), 100µl 1M NaCl (Cat# AM9760G Fisher Scientific, MA), 50ml nuclease-free water (Cat# AM9937 Thermo Fisher Scientific, MA), 25µl 1M spermine (Cat# 85590-5G, MilliporeSigma, MA), 10µl 1M spermidine (Cat# S2626-5G, MilliporeSigma, MA), 500µl proteinase inhibitor (PI; Cat# P9599-5ML MilliporeSigma, MA), 250µl BSA (Cat# B2518-100MG, MilliporeSigma, MA), 250µl SUPERase-In (Cat# AM2696, Thermo Fisher Scientific, MA)] and incubated for 10 min. After incubation, tissue was filtered through a 40µm filter (Cat# 43-57040-51, pluriSelect, Germany). We then centrifuged at 500 rcf for 5 min at 4°C. Supernatant was aspirated out, and nuclei were resuspended with NEB + 500µl 10% Triton (Cat# 93443-100ML, MilliporeSigma, MA) and no PI. We incubated for 15 min, filtered through a 40µm filter, and spun at 500 rcf for 5 min. We washed until the pellet was clear. We prepared the density gradient using the Density Buffer [DB; 120 mM Tris-Cl pH=8 (Cat# AM9855G, Fisher Scientific, MA), 150 mM KCl (Cat# AM9640G, Fisher Scientific, MA), 30 mM MgCl2, 35mL H_2_0 per 50mL] and filter sterilized it. We mixed 5 volumes of Optiprep (Cat# D1556-250ML, MilliporeSigma, MA) and the buffer volume to create a 50% stock. We also made a dilutant stock [400 mM Sucrose, 25 mM KCl, 5 mM MgCl_2_, 10 mM Tris-Cl pH 8, 28mL H_2_0 per 50mL] that we filter sterilized. A 45% solution was made by mixing 9ml 50% solution and 1ml dilutant, and a 15% solution by combining 1.5ml 50% stock and 3.5ml dilutant. We created a 45% solution and a 15% solution. We gently poured 2ml of the nuclei solution at the top of the density gradient and then spun the tubes at 1,500 rcf for 5 min with no breaks. After the spin, we pipette off nuclei and place them into a 15 ml tube with ∼ 6mL of NEB (with no triton or PI). We note that for vermiculite drought stressed nuclei, nuclei preparations were FACS purified. After counting the nuclei, the nuclei suspension was loaded onto microfluidic chips (10X Genomics) with HT-v3.1 chemistry to capture ∼20,000 nuclei/sample. Cells were barcoded with a Chromium X Controller (10X Genomics). mRNA was reverse transcribed, and Illumina libraries were constructed for sequencing with reagents from a 3’ Gene Expression HT-v3.1 kit (10X Genomics) according to the manufacturer’s instructions. cDNA and final library quality were assessed using Tape-Station High Sensitivity DNA Chip (Agilent, CA).

Each of the single-nuclei samples was processed twice, to get a higher number of single nuclei transcriptomes.

### Bulk RNA-sequencing and analysis

Sequencing was performed with a NovaSeq 6000 instrument (Illumina, CA). About 40 million reads were obtained for each sample. Raw reads were processed at the IGC bioinformatics core at Salk. Alignments were performed using OSA4 and mapped to the Arabidopsis genome (TAIR 10) using Tophat2 software with default settings. Mapped reads per library were counted using HTSeq software. Differentially expressed genes were quantified in two ways. Firstly, differentially expressed genes were identified using a spline regression model in splineTimeR v1.18.0, which were then sorted into time points using k-means clustering. Differential expression was also quantified at each time point individually using DESeq2 v1.30.0.^2^ For each pairwise comparison, genes with fewer than 32 total raw counts across all samples were discarded before normalization. Genes with an absolute log_2_foldchange > 1 and an FDR-corrected p-value ≤ 0.01 were pulled as significant. For functional enrichment, genes were queried for time-specific functional enrichment using over-representation analysis (ORA) in WebGestaltR v0.4.4.^3^ Differentially expressed genes in each pairwise comparison were queried against the biological process non-redundant ontology, and a significance threshold of FDR-corrected p-value ≤ 0.05 was used.

#### snRNA-seq analysis

For the snRNA-seq libraries, CellRanger (v.6.0.1) was used to perform sample-demultiplexing, barcode processing, and single-nuclei gene-UMI counting.^4^ Each experiment’s expression matrix was obtained by aligning to Arabidopsis transcriptome reference (TAIR 10) using CellRanger with default parameters. For initial quality-control filtering, aligned cell and transcript counts from each treatment (well-watered, drought, recovery, 2 replicates each) were processed by Seurat (Version 4.2).^5^ The data was filtered in the following two ways: (1) Pre-filtering each replicate by removing the low-quality and outlier cells containing a high abundance of chloroplast reads (>40% of total transcripts) and mitochondrial reads (>1% of total transcripts), a low abundance of detected genes (<300 detected genes) and a relatively high abundance of unique molecular identifiers (UMIs) (>10K for D0; >15K and >10K for two W15, respectively; >25K and >20K for two W0, respectively; >10K for R15) (Extended Data Table 3). (2) Identifying possible doublet in each replicate using the method SCDS. SCDS implements two complementary approaches to identify doublets: one is co-expression-based doublet scoring, and the other is binary classification-based doublet scoring. Additionally, they provide a hybrid score by combing these two approaches. SCDS showed relatively high detection accuracy and computational efficiency when benchmarking with other computational methods.^6^ We applied the hybrid scores for doublet estimation using R package scds() to identify likely doublet and then removed them from downstream integration. Expression data of cells passing these thresholds were log normalized with NormalizeData() function, and the top2K variable genes were identified with the FindVariableFeatures() function. Next, data from all conditions were integrated using Seurat’s reciprocal PCA (RPCA) and FindIntegrationAnchors() functions to identify integration features and correct for potential batch effects. The integrated data were then scaled with ScaleData() function. Principal component analysis (PCA) was carried out with RunPCA() function, and the top 30 principal components (PCs) were retained. Clusters were identified with the FindClusters() function using the shared nearest neighbor modularity optimization with a clustering resolution set to 0.8. Clusters with only one cell were removed. This resulted in 27 initial clusters with a total of 144,494 cells. We identified a median number of 1,500 genes, and 3,146 UMIs (representing unique transcripts), per nuclei. We detected between 25,064 – 26,342 genes in each sample. Cell-type identity of initial clusters was determined with canonical markers and by referring to public datasets, followed by sub-clustering each cluster using the same strategy described above.

Single cell differential gene expression analysis was conducted using Seurat FindMarker() and FindAllmarkers() functions with the default Wilcoxon Rank Sum test. Marker genes per cell cluster and differentially expressed genes (DEGs) among conditions were both identified by setting parameter min.pct as 0.1 and picked the top expressed markers ranking by average log_2_FC. Functional enrichment analysis was carried out by over-representation analysis (ORA) using the top100 markers ranked by average log_2_FC. Using the GO biological process as a reference, ORA was performed with WebGestalt.^3^ Pathways with FDR <0.05 were considered significantly enriched and visualized as plots with normalized enrichment scores. For recovery-/drought-specific DEGs, enriched known motifs were discovered with HOMER findmotifs.pl (http://homer.ucsd.edu/homer/), searching within 1000bp upstream to 1000kb downstream of their transcript start sites (TSSs). For the sub-clusters, recovery-enriched cell clusters were identified with >=50% cells from recovery experiment, followed with weighted gene co-expression network analysis (WGCNA) with R package hdWGCNA (https://smorabit.github.io/hdWGCNA/). Co-expression networks were constructed for each initial cell cluster with top 20 hub genes. By checking the average expression of hub genes in sub-clusters, the recovery-specific networks were picked as those with higher expression in recovery-enriched cell clusters.

### Statistical analysis

bulk RNA-seq data, statistical analysis was performed in R using a mixed linear model function (lmer) from the package lme4 unless otherwise described. Standard errors were calculated from variance and covariance values after model fitting. The Benjamini-Hochberg method was applied for correcting of multiple testing in figures showing all pairwise comparisons of the mean estimates. For bacterial colonies count, merged two independent experiments of plants grown on plates, a total of n=12 per treatment. Significance values for log10(CFU) on the plates were calculated with a two-way ANOVA of treatment and batch followed by a Tukey test. P-values are FDR-corrected. For bacterial colonies count, soil grown plants experiment, significance values for log10(CFU) were calculated with a two-way student’s t-test.

## Data availability

Data is available here: GSE220278 (https://www.ncbi.nlm.nih.gov/geo/query/acc.cgi?acc=GSE220278). Currently open only for reviewers.

## Code availability

The code used to analyze both bulk and single-nuclei RNA-Seq data is available at: https://github.com/NatanellaIE/DroughtRecovery.

## Acknowledgments

We want to thank T. Nolan for insightful comments to this manuscript. This research was supported by a Postdoctoral Award No. FI-601-2020 from the United States–Israel Binational Agricultural Research and Development Fund. N.I.-E. is a research fellow at the George E. Hewitt Foundation for Medical Research and an Awardee of the Weizmann Institute of Science – Israel National Postdoctoral Award Program for Advancing Women in Science and the Women’s Postdoctoral Career Development Award. J.R.E. is an Investigator of the Howard Hughes Medical Institute.

## Author contributions

Conceptualization – N.I-E

Data curation – N.I-E, R.G-C, J.S, B.J, T.Z-K, A.Y

Formal analysis – N.I-E, K.L, J.Y, J.R.N, T.Z-K

Funding acquisition - N.I-E, J.R.E

Investigation - N.I-E, R.G-C

Methodology - N.I-E, T.L, J.S, S.B, Y.Z

Project administration - N.I-E

Resources – J.R.E

Writing – original draft - N.I-E, J.R.E.

## Declaration of interests

The authors declare no competing interests.

## Notes

https://github.com/NatanellaIE/DroughtRecovery

### Competing Interest Statement

The authors have declared no competing interest.

### Summary of Updates

This revised version of the manuscript includes the following updates: We have resequenced our single-nuclei libraries to achieve greater depth. Our updated single-cell dataset now comprises over 144,000 nuclei, with ^1,500 genes per nucleus. Reanalysis of this expanded dataset has led to several new insights: First, we observed a rapid recovery-induced transcriptional reprogramming specifically within dividing cells. Next, we identified a distinct cell state induced by recovery, which forms independently across various cell types. Additionally, to assess whether the DRII response is specific to Arabidopsis or more broadly applicable, we investigated whether short-term recovery from moderate drought enhances pathogen resistance in both wild and domesticated tomato. Our findings suggest that the DRII response is conserved in tomato and was conserved during the domestication process.

