## Extended Data Figures for "Stress Recovery Triggers Rapid Transcriptional Reprogramming and Activation of Immunity in Plants"

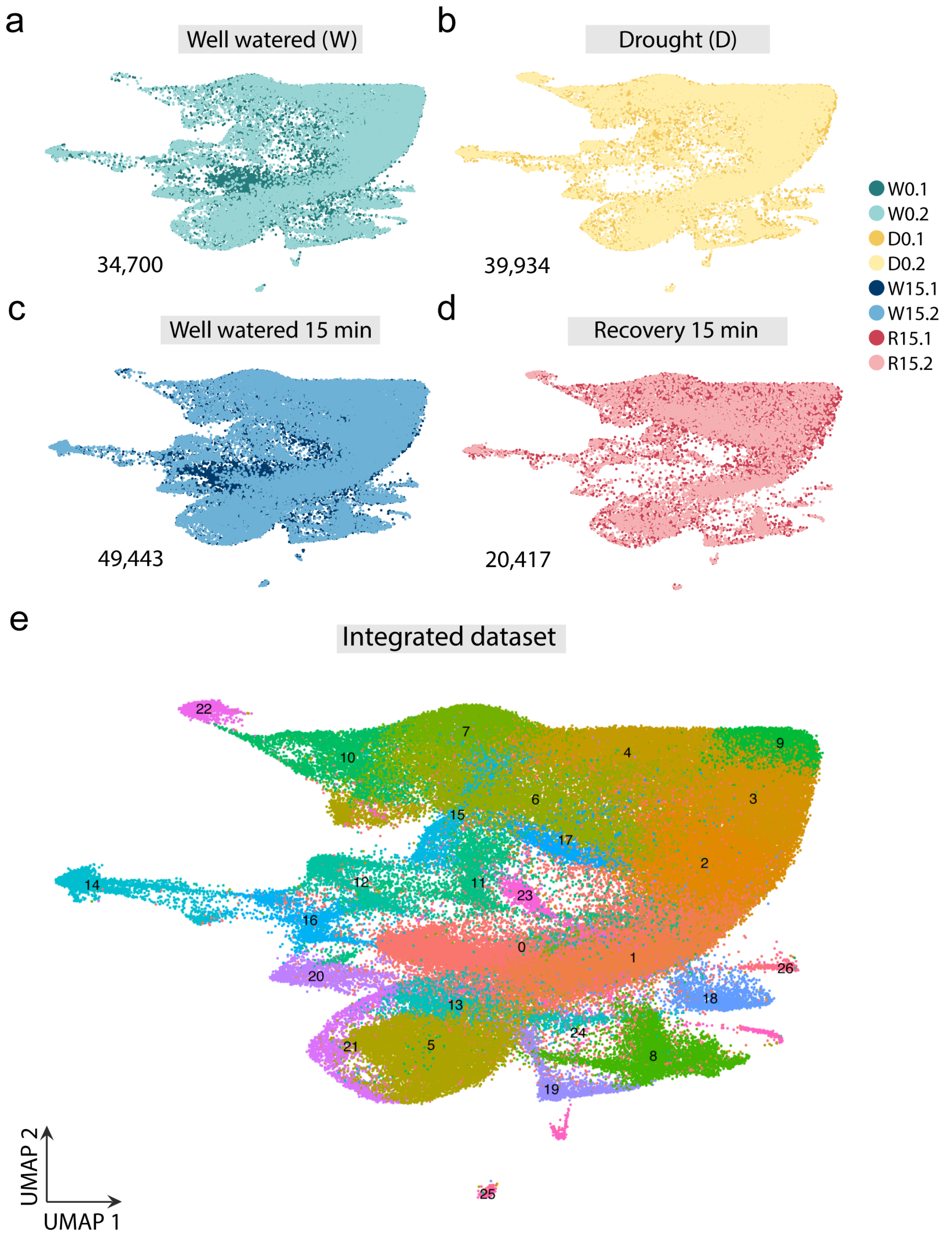


**Extended Data Figure 1. The biological replicates and different conditions integrate into similar cell identities.** UMAP projection of 2 independent samples of **a,** Well-watered plants, **b,** Drought treated plants, **c,** Well watered after 15 mins of additional hydration and **d,** Drought treated plants after rehydration for 15 mins. **e,** Seurat clusters of the integrated dataset combining all independent samples.

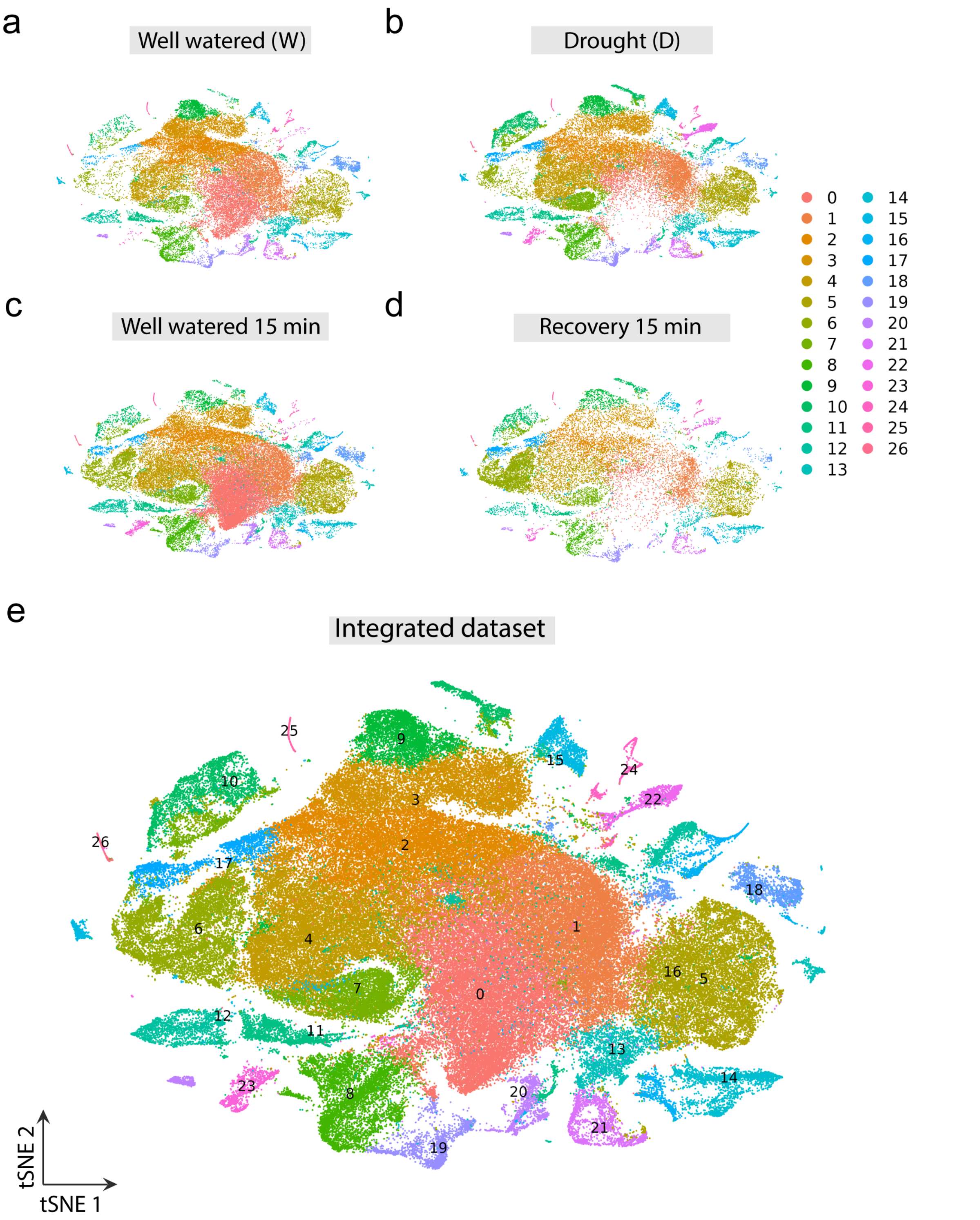


**Extended Data Figure 2. The biological replicates and different conditions integrate into similar cell identities using tSNE.**

tSNE projection of 2 independent samples of **a,** Well-watered plants, **b,** Drought treated plants, **c,** Well watered after 15 mins of additional hydration and **d,** Drought treated plants after rehydration for 15 mins. **e,** Seurat clusters of the integrated dataset combining all independent samples.

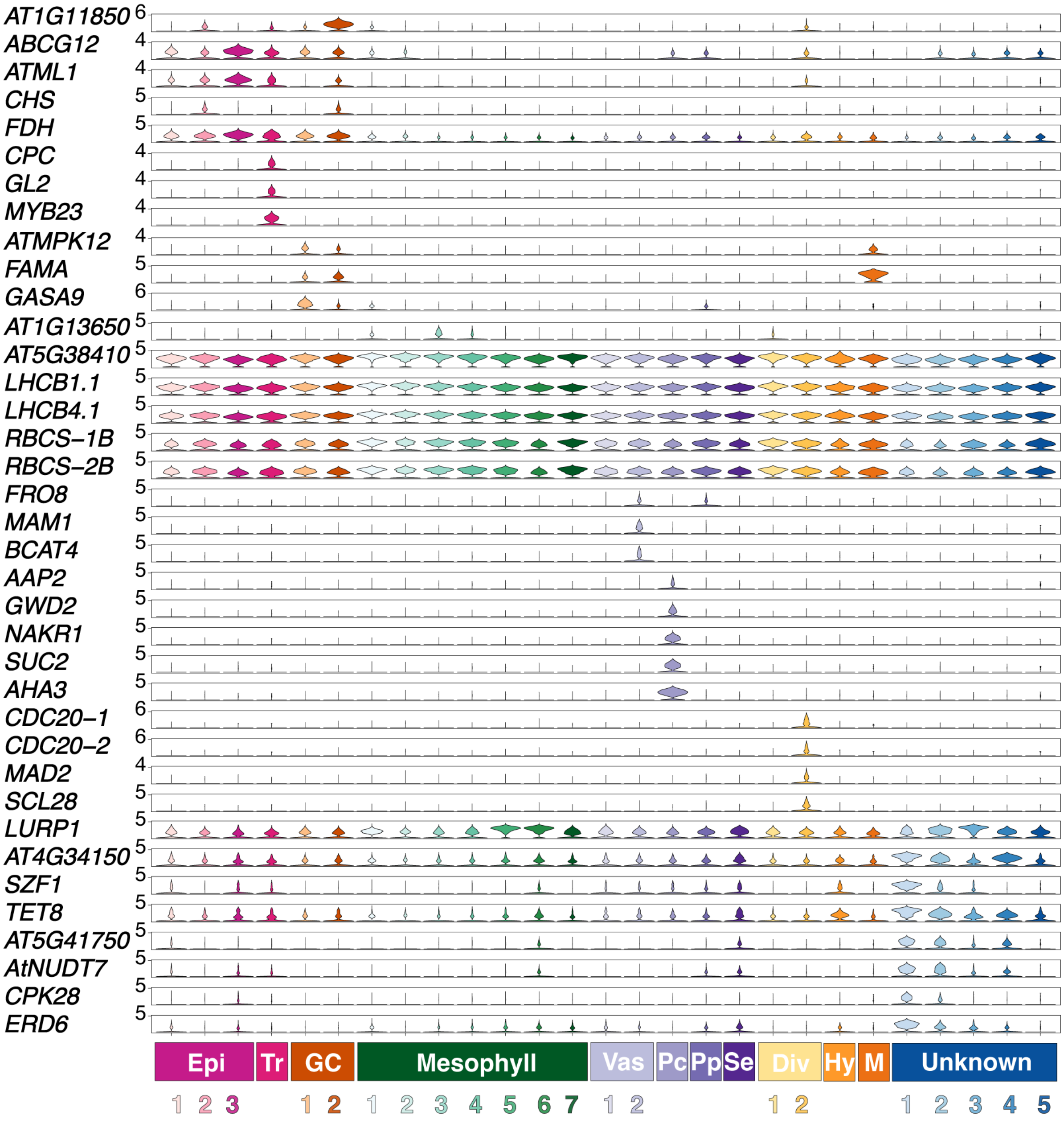


**Extended Data Figure 3. Expression of literature tissue- and cell-type specific genes.** Violin plots presenting the expression of genes in different clusters of the integrated dataset revealing 27 cell identities encompassing the major leaf cell types. Cell type identities: Epi = Epidermal, GC = Guard Cell, Tr = Trichome, Hy = Hydathode, M = Myrosin, Mes = Mesophyll, Pp = Phloem parenchyma, Pc = Phloem companion, Se = Sieve element, Div = Dividing, and five unknown clusters- Unknown 1-5.

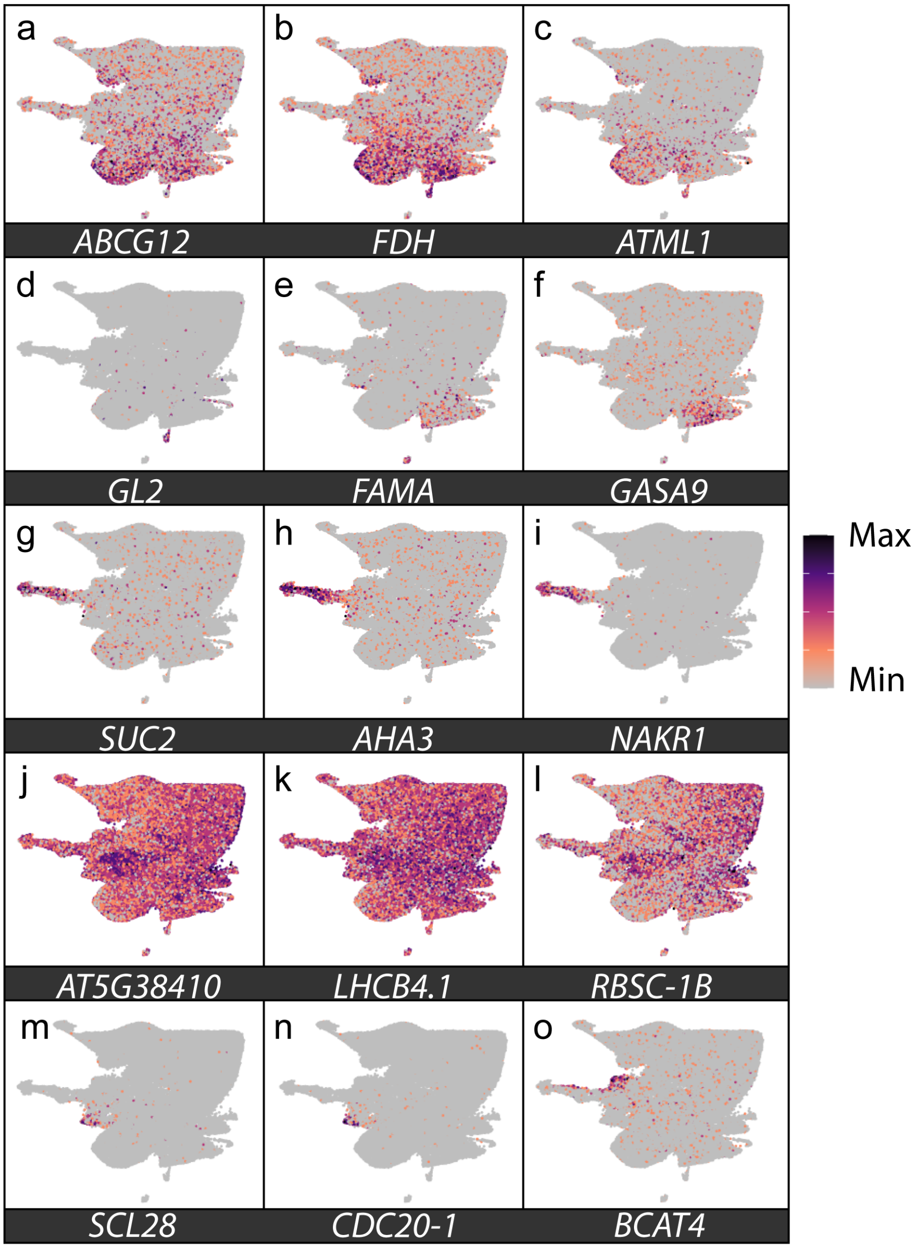


**Extended Data Figure 4. Example literature tissue- and cell-type specific marker genes projected on our integrated UMAP. a,** *ARABIDOPSIS THALIANA WHITE-BROWN COMPLEX 12* (*ABCG12*), encodes an ABC transporter involved in cuticular wax biosynthesis. **b,** *FIDDLEHEAD* (*FDH*), epidermis-specific, encodes KCS10, a putative 3-ketoacyl-CoA synthase. probably involved in the synthesis of long-chain lipids found in the cuticle. **c,** *MERISTEM LAYER 1* (*ATML1*), epidermis specific, encodes a homeobox protein similar to GL2. **d,** *GLABRA 2* (*GL2*) a homeodomain protein affects epidermal cell identity including trichomes. **e,** *FAMA* (*FMA*), Encodes a basic helix-loop-helix transcription factor whose activity is required to promote differentiation of stomatal guard cells and to halt proliferative divisions in their immediate precursors. **f,** *GIBBERELLIC ACID STIMULATED ARABIDOPSIS 9* (*GASA9*). **g,** *SUCROSE-PROTON SYMPORTER 2* (*SUC2*), Encodes for a high-affinity transporter essential for phloem loading and long-distance transport. A major sucrose transporter. **h,** *Arabidopsis H(+)-ATPase isoform 3* (*AHA3*)**. i,** *SODIUM POTASSIUM ROOT DEFECTIVE 1* (*NaKR1*) encodes a phloem mobile metal binding protein necessary for phloem function and root meristem maintenance. **j,** *RUBISCO SMALL SUBUNIT 3B* (*RBCS3B*), encodes a member of the Rubisco small subunit (RBCS) multigene family. **k,** *LIGHT HARVESTING COMPLEX PHOTOSYSTEM II* (*LHCB4.1*). **l,** *RUBISCO SMALL SUBUNIT 1B* (*RBCS1B*)**. m,** *SCARECROW-LIKE 28* (*SCL28*) transcription factor belonging to the GRAS family which controls the mitotic cell cycle and division plane orientation. **n,** *CELL DIVISION CYCLE 20.1* (*CDC20.1*) Encodes a CDC20 protein that interacts with APC subunits, components of the mitochondrial checkpoint complex and mitotic cyclin substrates and is indispensable for normal plant development and fertility. **o,** *BRANCHED-CHAIN AMINOTRANSFERASE 4* (*BCAT4*) belongs to the branched-chain amino acid aminotransferase gene family. Encodes a methionine-oxo-acid transaminase.

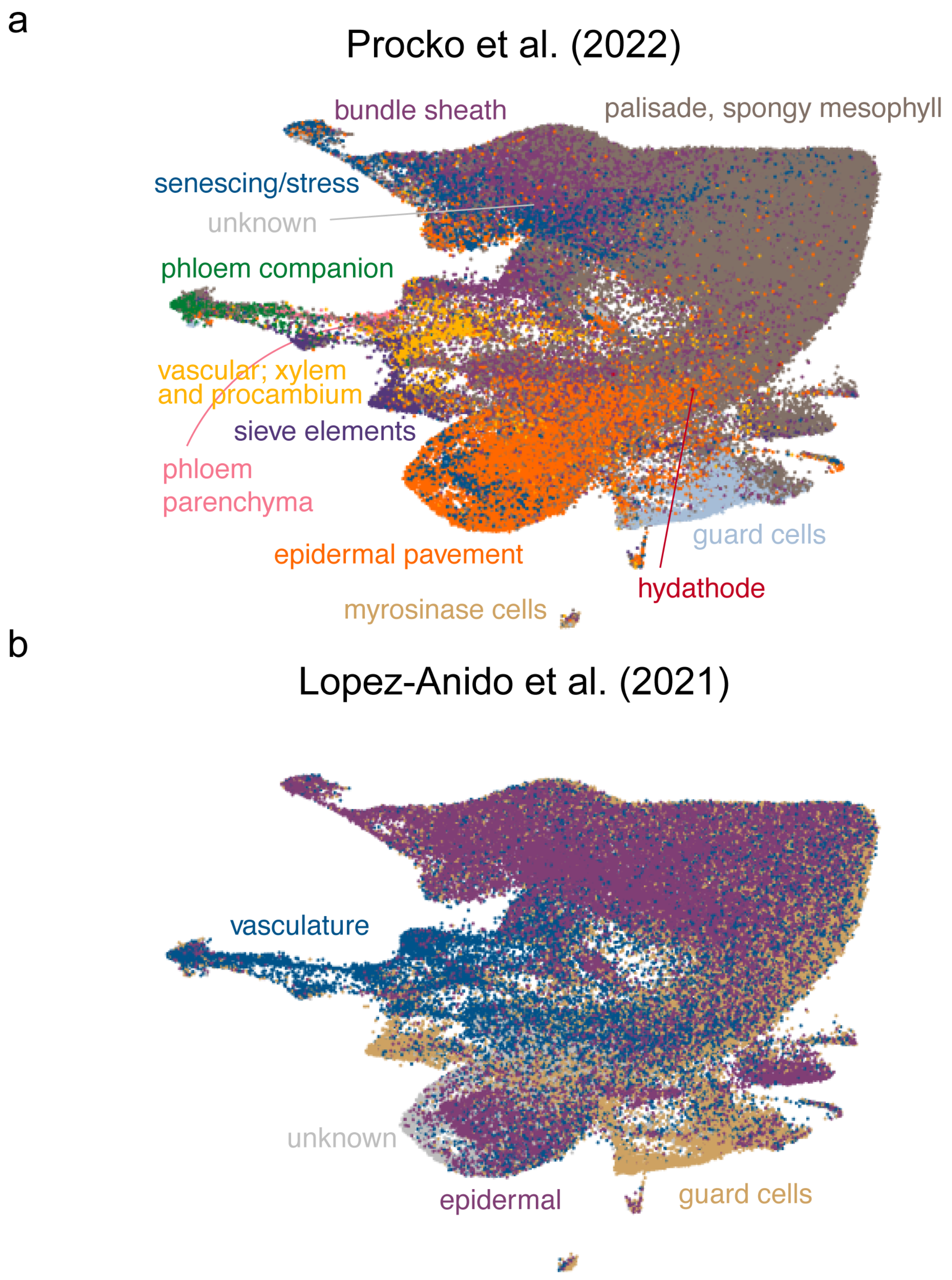


**Extended Data Figure 5. Projection of available annotated Arabidopsis leaf single-cell datasets onto our single-nuclei RNA-seq clusters improves cluster annotation. a,** Projection of single-cell RNA-seq annotation information from Procko et al., 2022 onto our Arabidopsis rosette single-nucleus dataset. **b,** Projection of single-cell RNA-seq data from Lopez-Anido et al., 2021.

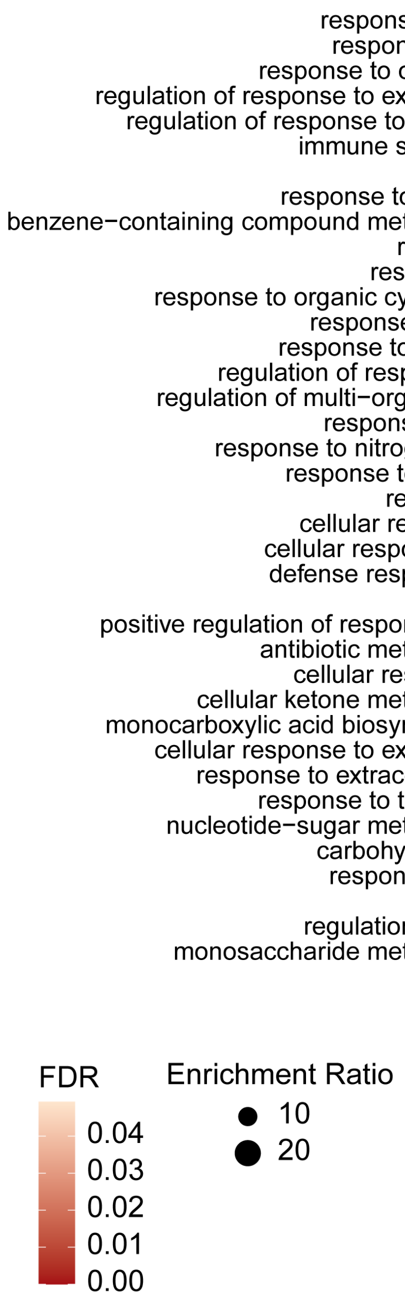


**Extended Data Figure 6. Gene Ontology (GO) enrichment analysis of the 5 unidentified clusters.** GO enrichment analysis performed on five clusters that were not previously identified or annotated. Unknown 5 shows enrichments in nucleotide-sugar metabolic processes which may indicate potential alterations in glycosylation patterns, cell wall biosynthesis, and other cellular processes dependent on nucleotide-sugar substrates (Bar-Peled and O'Neill, 2011). Enrichment in carbohydrate transport which suggests an upregulation or downregulation of genes involved in the movement of carbohydrates across cellular membranes, indicative of alterations in energy metabolism, and nutrient uptake (Lalonde et al., 2004). Additionally, we observed enrichment in response to metal ion which indicates activation of genes involved in metal ion homeostasis, detoxification, and stress responses (Cobbett and Goldsbrough, 2002).

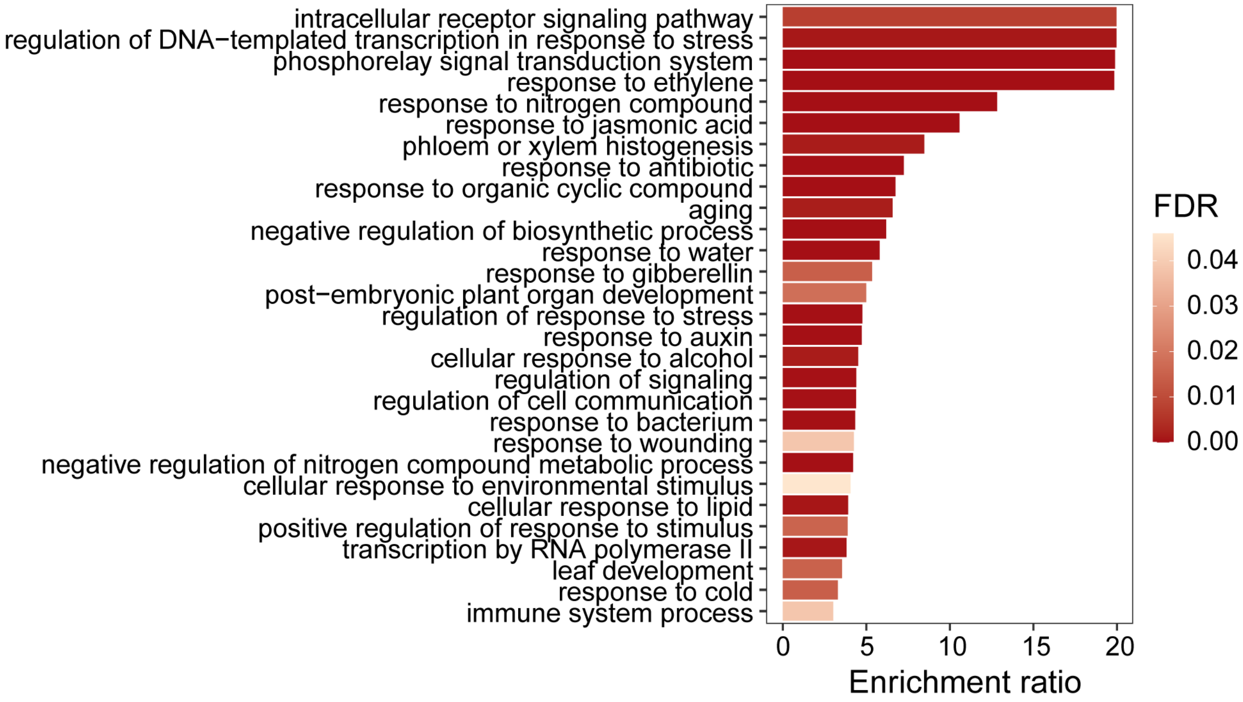


**Extended Data Figure 7. Gene Ontology (GO) enrichment analysis of transcription factors globally induced by rehydration across cell-types.** GO enrichment analysis performed on 136 TFs that were upregulated specifically by rehydration, in all cell types.

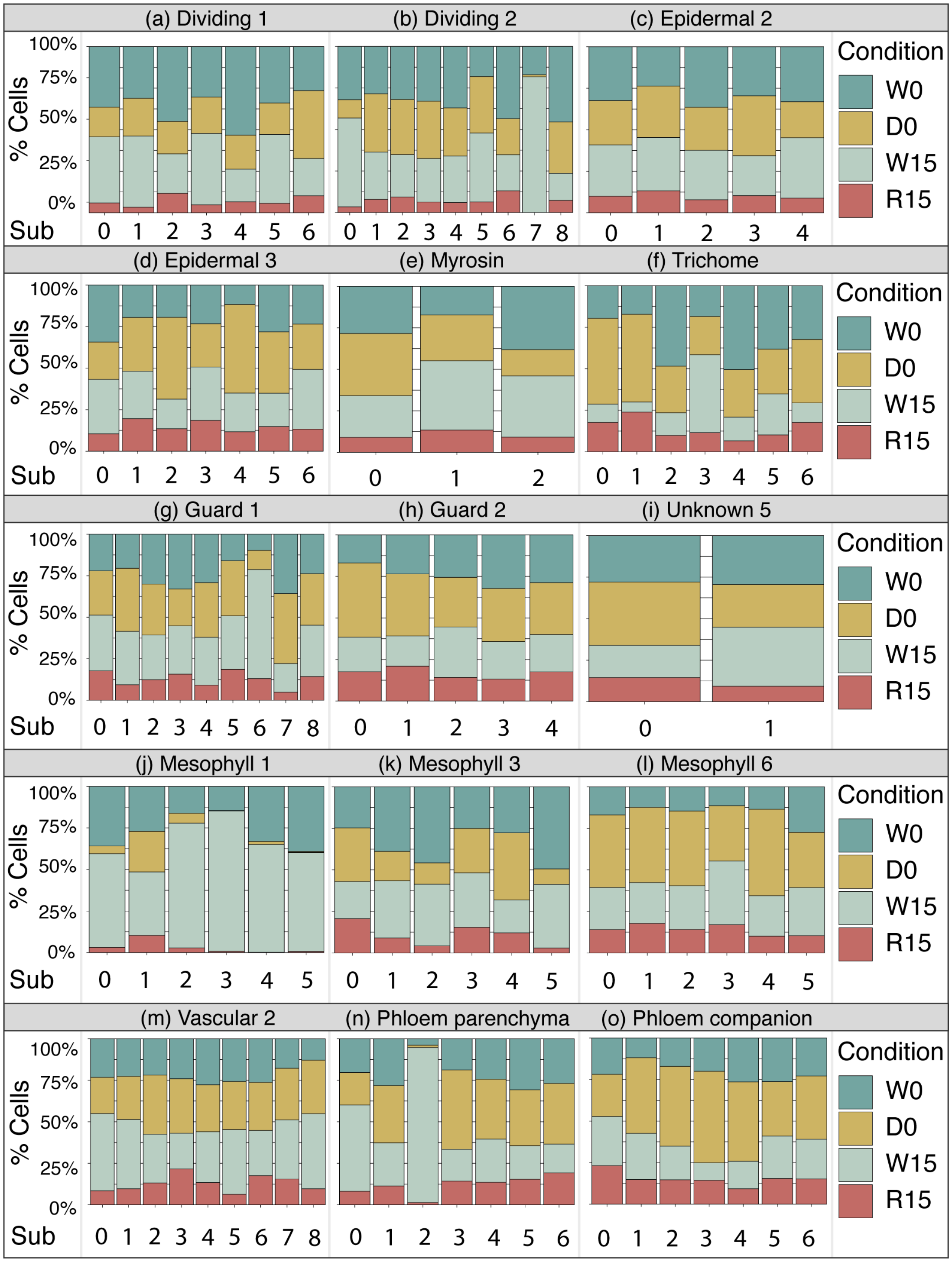


**Extended Data Figure 8. The distribution of cells by condition in subclusters of major cell identity clusters.** The bar plots present cell percentage in **a,** Dividing cells - 1 **b,** Dividing cells - 2 **c,** Epidermal cells - 2 **d,** Epidermal cells – 3 **e,** Myrosin cells **e,** Trichome cells **g,** Guard cells 1 **h,** Guard cells 2 **i,** Unknown 5 **j,** Mesophyll 1 **k,** Mesophyll 3 **l,** Mesophyll 6 **m,** Vascular 2 **n,** Phloem parenchyma **o,** Phloem companion**.**

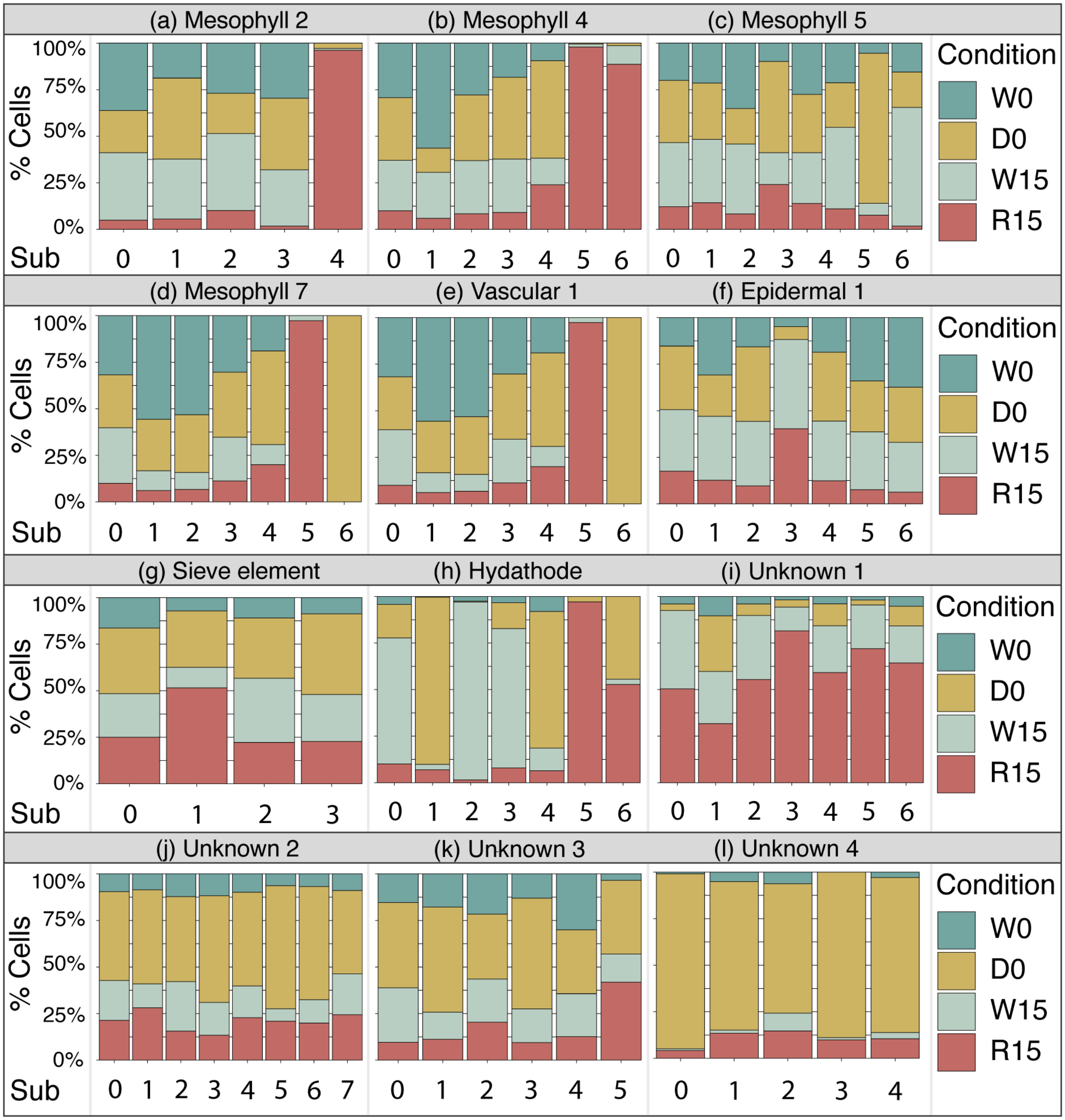


**Extended Data Figure 9. The distribution of cells by condition in subclusters of major cell identity clusters.** The bar plots present cell percentage in **a,** Mesophyll 2 **b,** Mesophyll 4 **c,** Mesophyll 5 **d,** Mesophyll 7 **e,** Vascular 1 **f,** Epidermal 1 **g,** Sieve element **h,** Hydathode **i,** Unknown 1 **j,** Unknown 2 **k,** Unknown 3 **l,** Unknown 4.

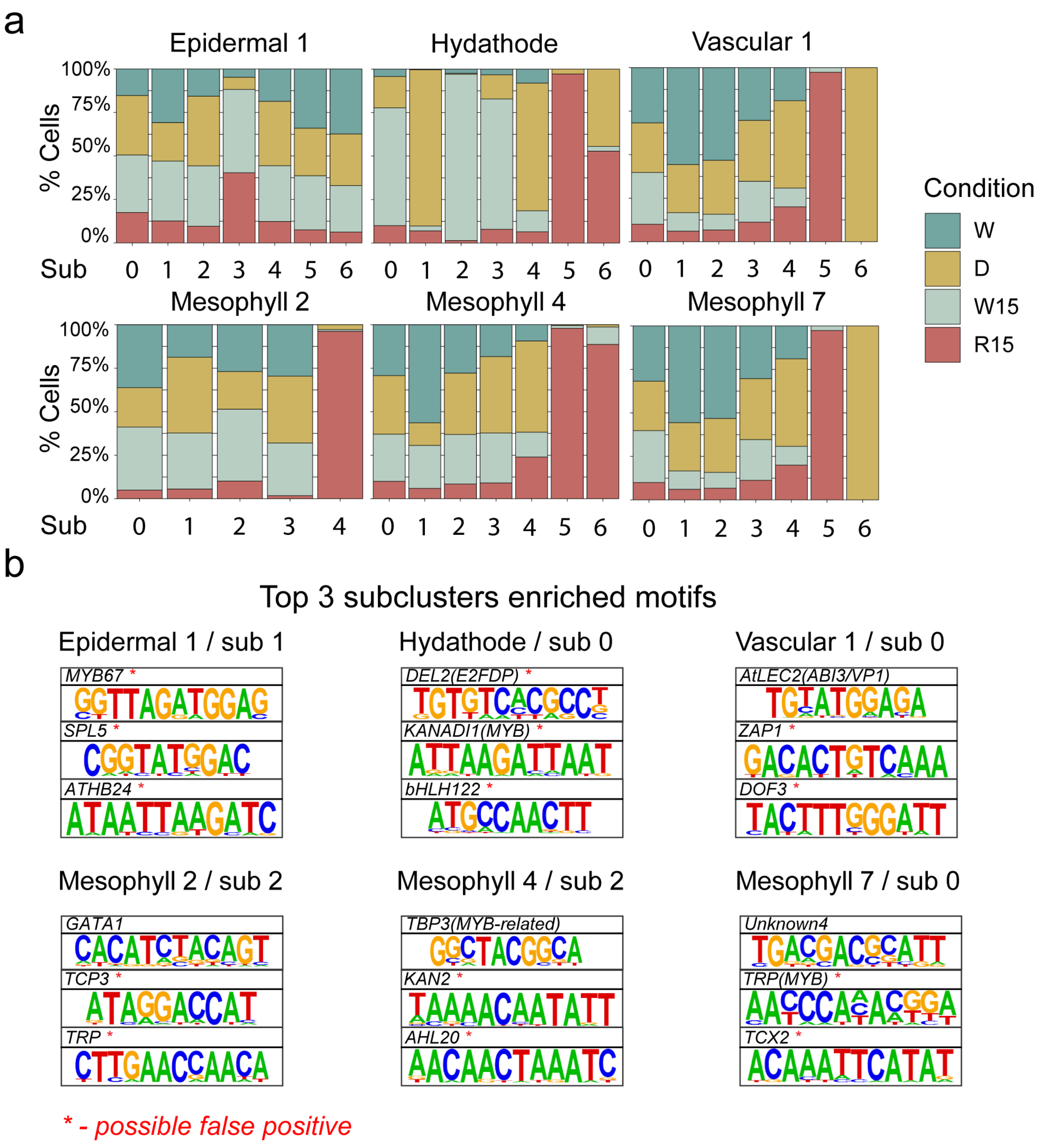


**Extended Data Figure 10. Motif enrichment analysis of subclusters with equal representation of cells by condition do not show enrichment in CAMTA binding motifs. a,** Bar plots present cell percentage in cell identities. The subcluster we did the analysis on were: Epidermal 1 – subcluster 1, Hydathode – subcluster 0, Vascular 1 – subcluster 0, Mesophyll 2 – subcluster 2, Mesophyll 4 – subcluster 2, and Mesophyll 7 – subcluster 0. **b,** *de novo* Motif enrichment analysis for each recovery enriched subcluster. The gene name above the motif is the predicated TF binding the enriched motif. The analysis was performed with HOMER, using the top 100 marker from each cluster with FDR<0.05.

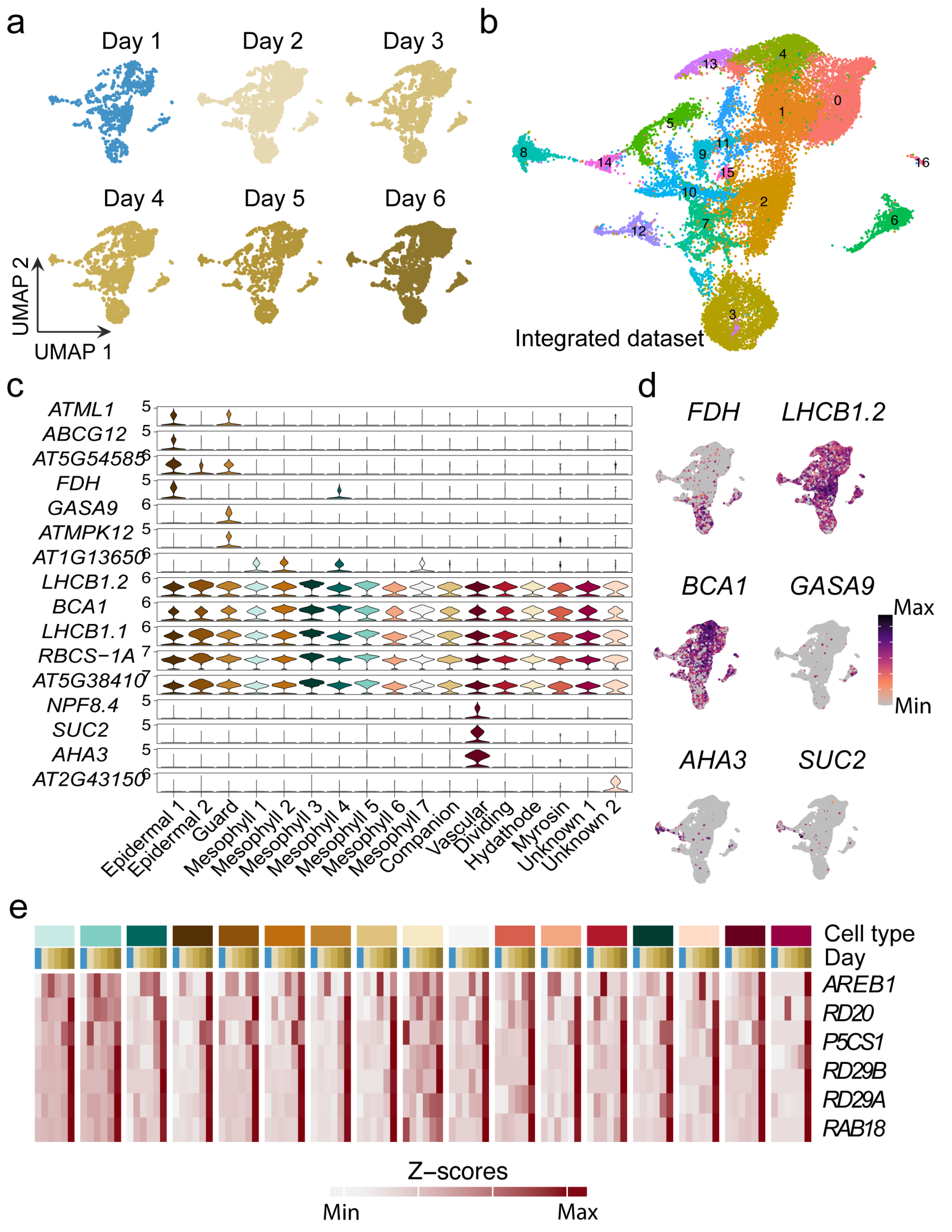


**Extended Data Figure 11. snRNA-seq of plant in early stages of dehydration.**
**a,** UMAP projection of 5 days of dehydration from a well-watered state (Day 1). **b,** Seurat clusters of the integrated dataset combining all samples. **c,** Violin plots presenting the expression of genes in different clusters of the integrated dataset. **d,** Example literature tissue- and cell-type specific marker genes projected on our integrated UMAP. **e,** The expression levels of drought inducible genes in the drought snRNA-seq dataset.

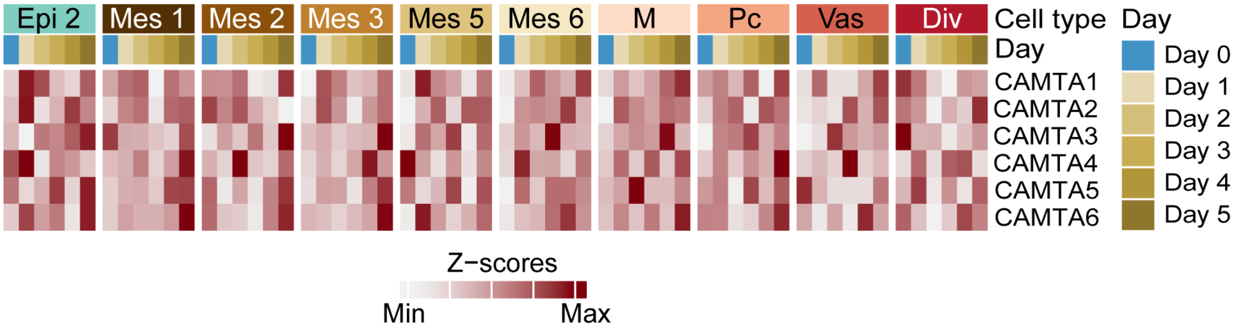


**Extended Data Figure 12. The expression of CAMTA TFs in early drought stages.** Z-score representation of the expression levels of the predicted CAMTA TFs during early stages of plants dehydration in the drought dataset.

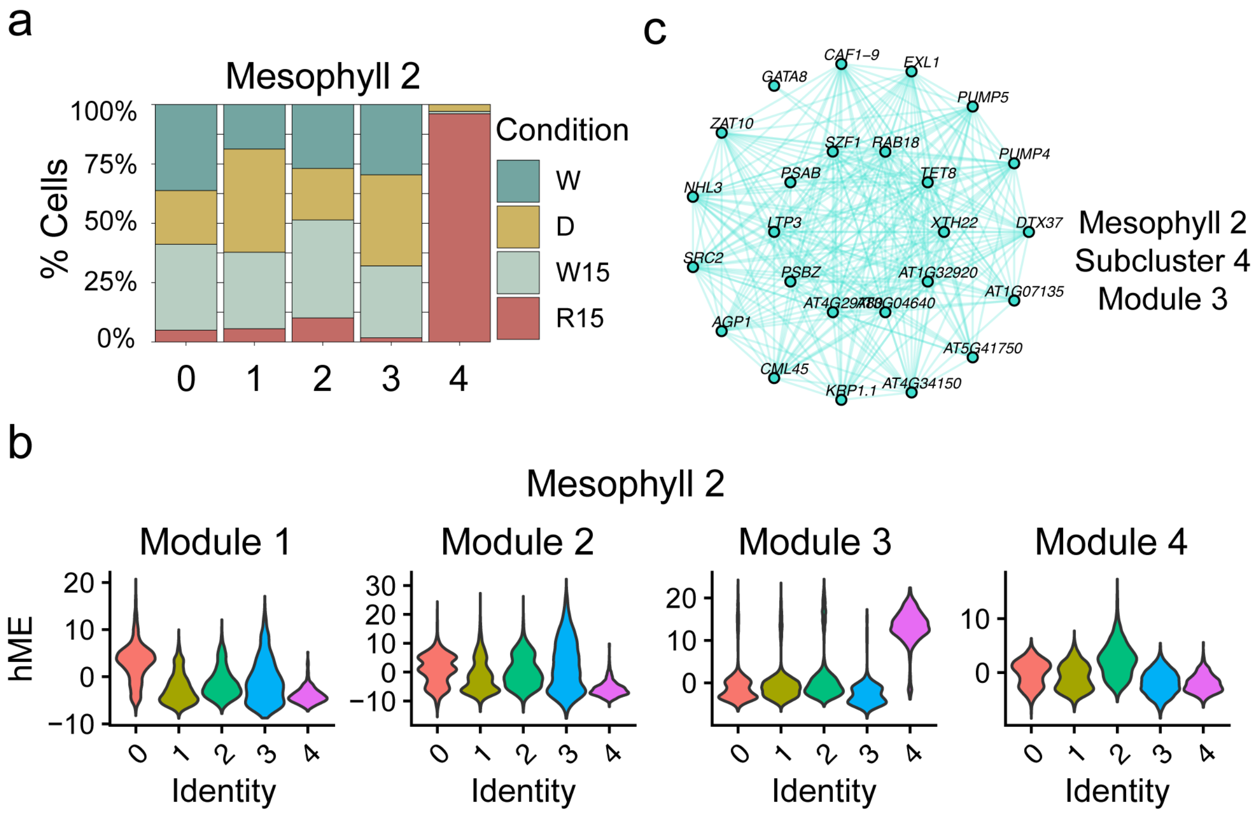


**Extended Data Figure 13. Gene module identification in putative RcS.** An example of the gene network analysis preformed to identify shared networks and hub genes between RcS activated in different cell types. In this example, **a,** shown subclustering of cluster “Mesophyll 2”, where subcluster 4 is enriched, almost exclusively in cells from the onset of recovery. **b,** Gene modules identified and there aboundance in each subcluster. Module 3 is enriched in subcluster 4. **c,** Top 25 hub genes of module 3. Each cell type subclusters enriched in recovery cells (over 50% of all cells in the subcluster), were used for the analysis, and hub genes were used for downstream analyses.

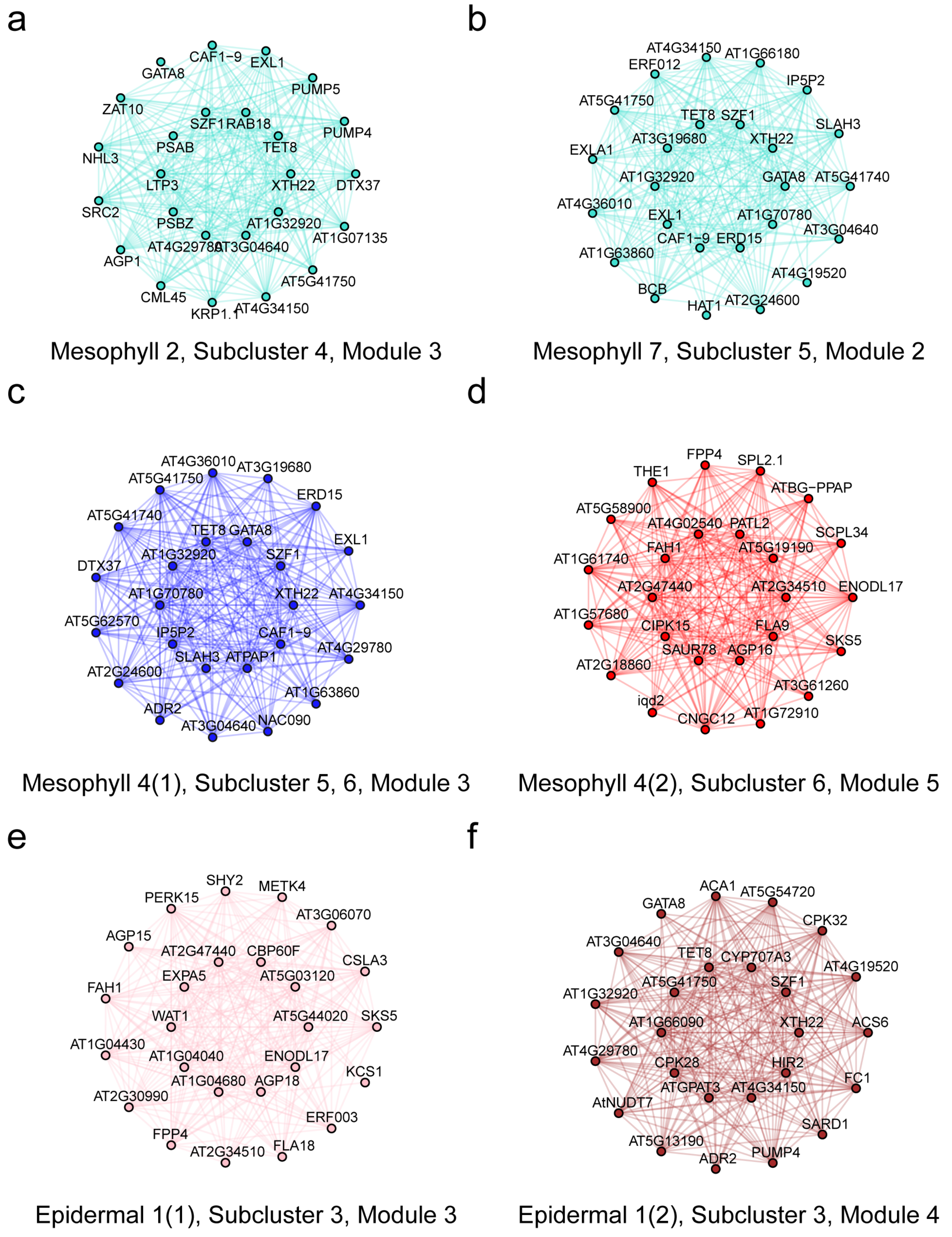


**Extended Data Figure 14. Top 25 hub genes of the gene modules enriched in each RcS**. It should be noted that if there are two subclusters enriched with recovery cells in the same cell type cluster, the algorithm will not allow them to have overlapping hub genes.

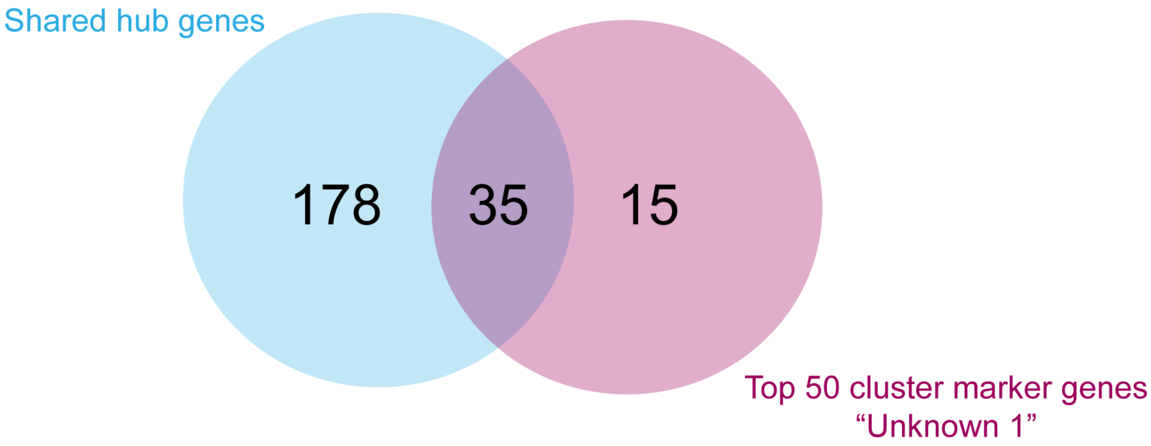


**Extended Data Figure 15. The overlap between RcS shared hub genes and the top 50 cluster marker genes of recovery enriched unknown 1 cluster.** Venn diagram exhibits 70% of the top 50 unknown 1 cluster markers are found as hub genes of RcS populations.

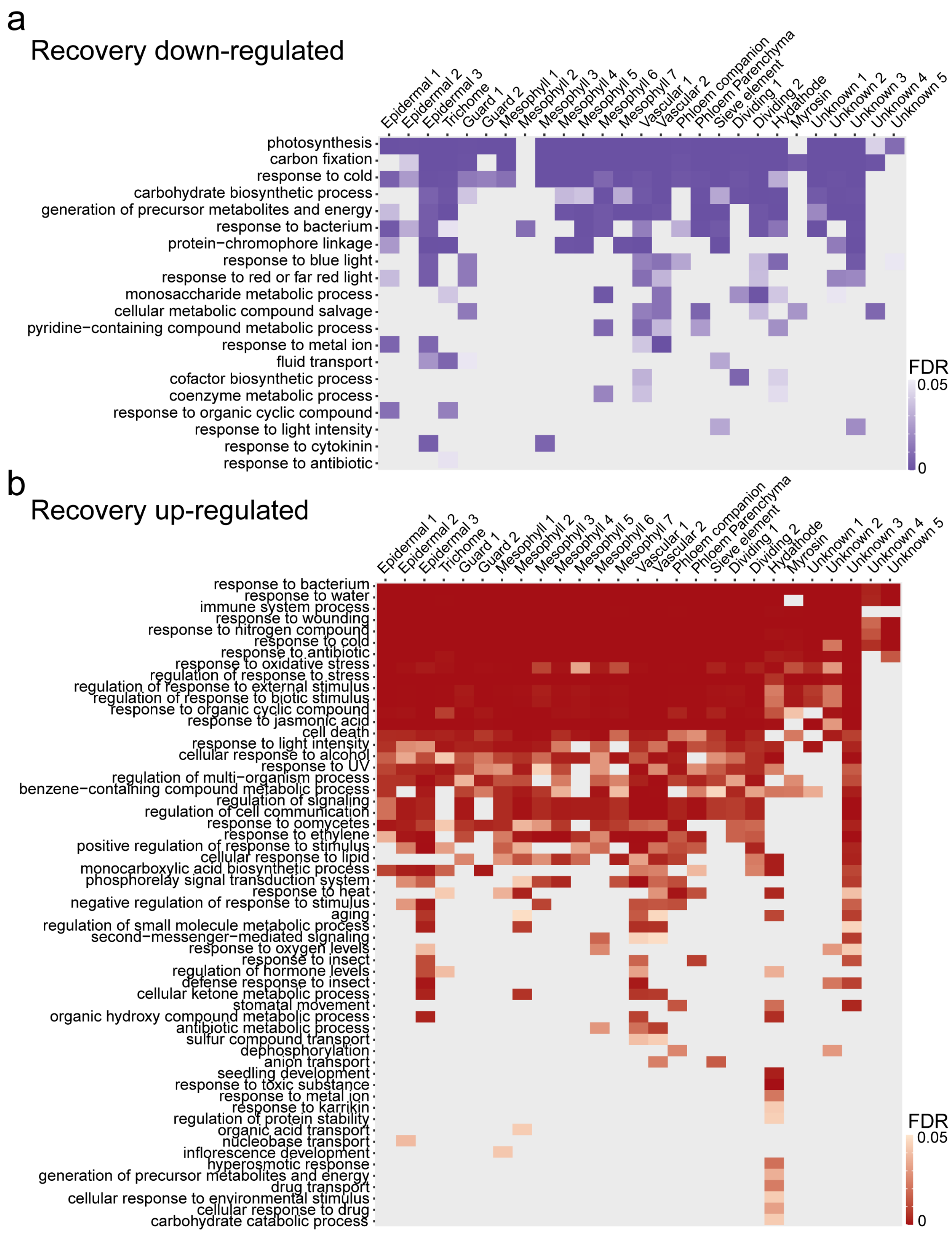


**Extended Data Figure 16. GO enrichment analysis of the genes induced or repressed by recovery.** GO enrichment analysis for genes DE down **a,** or **b,** up by recovery compared to drought samples as well as well-watered 15 minutes control.

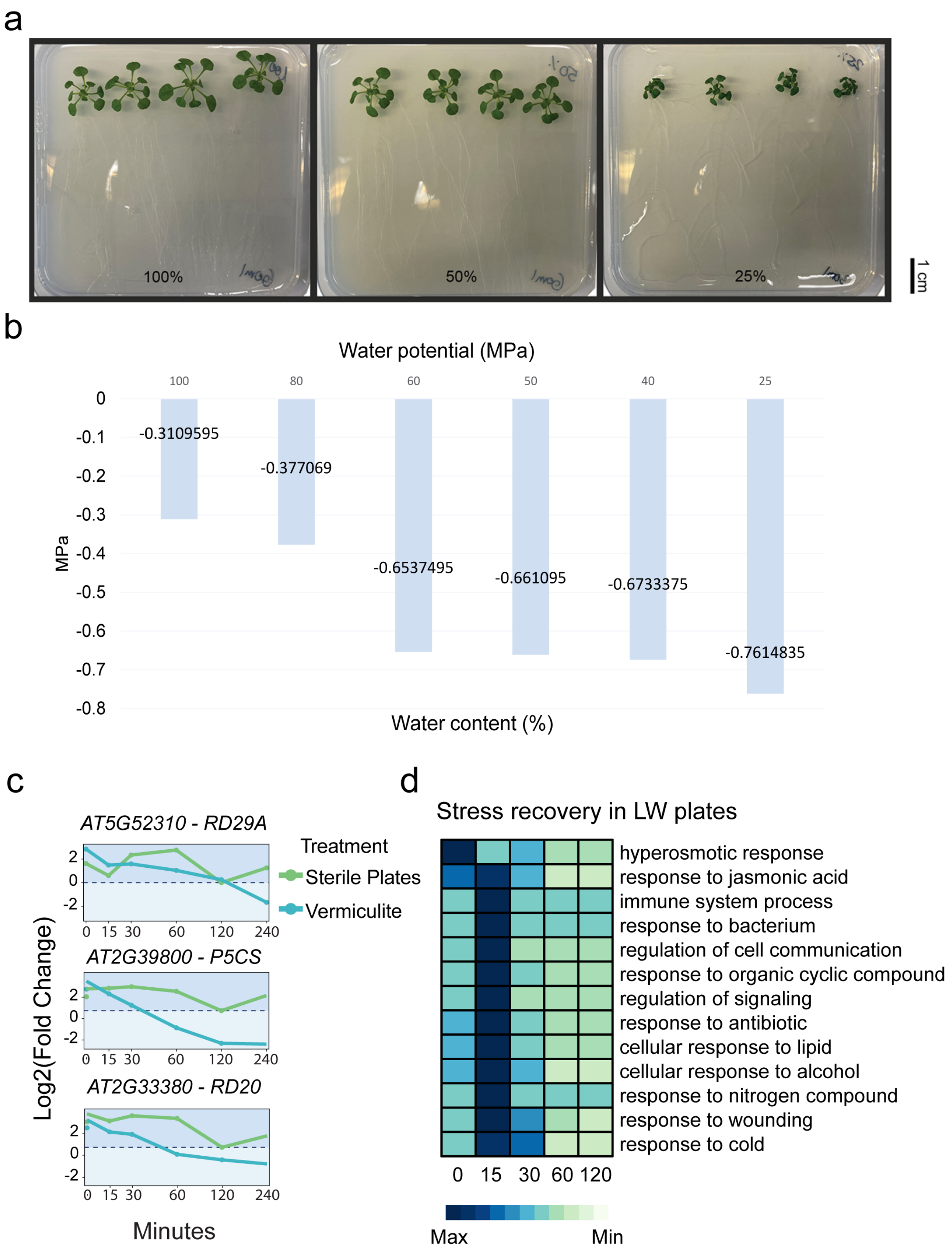


**Extended Data Figure 17. LW plate system for simulating drought stress in a sterile environment. a,** plant phenotype on low water (LW) plates. **b,** water potential values of LW plates. **c,** Expression of drought marker genes: *RD29A*, *P5CS* and *RD20*. **d,** GO enrichment analysis for genes up-regulated by moderate stress (time 0) and recovery in the low water (LW) sterile plates.
